# Cyclin E/CDK2 and feedback from soluble histone protein regulate the S phase burst of histone biosynthesis

**DOI:** 10.1101/2023.03.17.533218

**Authors:** Claire Armstrong, Victor J. Passanisi, Humza M. Ashraf, Sabrina L. Spencer

## Abstract

Faithful DNA replication requires that cells fine-tune their histone pool in coordination with cell-cycle progression. Replication-dependent histone biosynthesis is initiated at a low level upon cell-cycle commitment, followed by a burst at the G1/S transition, but it remains unclear how exactly the cell regulates this change in histone biosynthesis as DNA replication begins. Here, we use single-cell timelapse imaging to elucidate the mechanisms by which cells modulate histone production during different phases of the cell cycle. We find that CDK2-mediated phosphorylation of NPAT at the Restriction Point triggers histone transcription, which results in a burst of histone mRNA precisely at the G1/S phase boundary. Excess soluble histone protein further modulates histone abundance by promoting the degradation of histone mRNA for the duration of S phase. Thus, cells regulate their histone production in strict coordination with cell-cycle progression by two distinct mechanisms acting in concert.

## Introduction

Maintaining genomic stability during DNA replication requires that cells double their histone proteins in coordination with the doubling of the genome beginning at the start of S phase when DNA replication is initiated. Cells must rapidly produce approximately 400 million replication-dependent histone proteins, herein referred to as histone proteins, in as little as 8 hr to accommodate genome doubling in S phase (Duronio and Marzluff, 2017). Our recent work established a temporal decoupling of the initiation of low levels of histone biosynthesis at cell-cycle commitment and a burst of histone production 5 hr later at the G1/S boundary (Armstrong and Spencer, 2021; Moser et al., 2018; Spencer et al., 2013). Thus, new questions have arisen as to how cells regulate this transition from low to high levels of histone biosynthesis at the start of S phase.

Metazoans have evolved a tightly regulated system for histone biosynthesis. In humans, histone genes are found in two evolutionarily conserved clusters at chromosomes 1 (HIST2) and 6 (HIST1) around which a nuclear body forms, known as the histone locus body (HLB) (Duronio and Marzluff, 2017; Marzluff et al., 2002; Tatomer et al., 2016). HLB formation is dependent on the scaffolding protein and histone transcription factor, nuclear protein at the ATM locus (NPAT) (Tatomer *et al*., 2016). Several other HLB specific factors are recruited upon HLB formation, including FLICE-associated huge protein (FLASH), which then recruits U7 snRNA-associated Sm-like protein (Lsm11) for 3’ end processing of histone mRNA (Skrajna et al., 2017; Tatomer *et al*., 2016; Yang et al., 2009; Yang et al., 2013). Once transcribed, histone mRNA is not polyadenylated at the 3’ end, but instead ends in a 3’ stem loop that requires binding to the stem loop binding protein (SLBP) for stabilization (Figure 1A) (Marzluff and Koreski, 2017; Sullivan et al., 2009; Whitfield et al., 2000). In the absence of SLBP, histone mRNA will be rapidly degraded by histone specific exonuclease 3’hExo (Koseoglu et al., 2008; Marzluff and Koreski, 2017; Meaux et al., 2018; Mendiratta et al., 2019; Mullen and Marzluff, 2008; Zheng et al., 2003).

**Figure 1.**
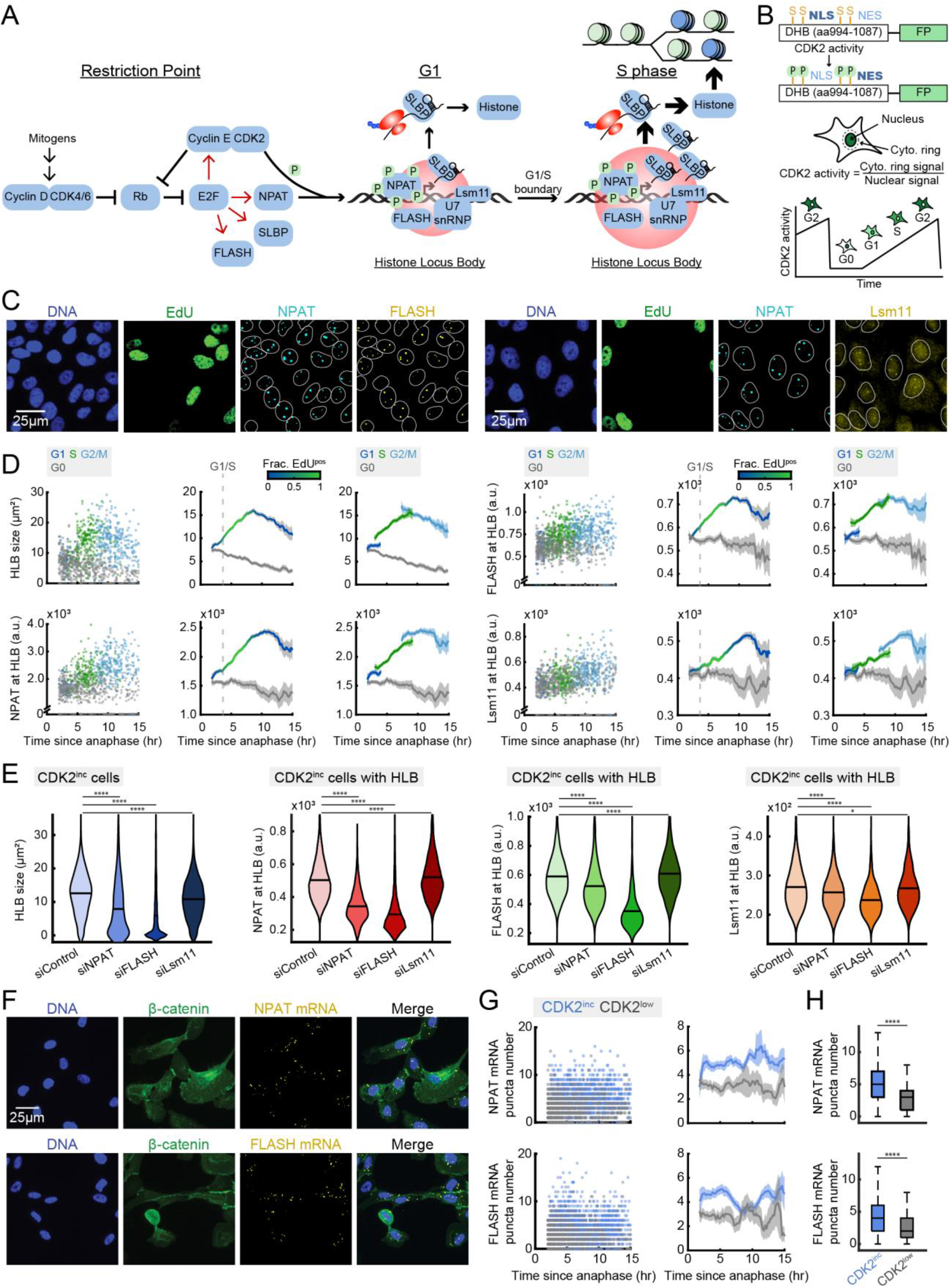
NPAT and FLASH are required for HLB formation. (A) Schematic of histone biosynthesis relative to the Restriction Point and cell-cycle progression. (B) Schematic of the CDK2 sensor and live-cell tracking (Spencer *et al*., 2013). (C) Representative images of cells following timelapse imaging, stained for DNA content, EdU, NPAT, and either FLASH or Lsm11. (D) Column 1: Raw single-cell data by cell-cycle phase. Column 2: Average of the total HLB size per cell or protein signal at the HLB and 95% confidence intervals as a function of time since anaphase for CDK2^inc^ and CDK2^low^ populations, with G1/S transition as the time when 50% of cells are EdU^pos^. Column 3: CDK2^inc^ population from column 2 segmented in to G1, S, and G2/M (see also Figure S1). (E) Violin plots of average HLB size for all CDK2^inc^ cells and average intensity for CDK2^inc^ cells with HLB puncta for NPAT, FLASH, and Lsm11 protein following 48 hr treatment with siControl, siNPAT, siFLASH, or siLsm11. (F) Representative images of cells following timelapse imaging and staining for DNA, β-catenin for cytoplasmic segmentation, and either NPAT or FLASH mRNA. (G) Column 1: Raw single-cell data for CDK2^inc^ and CDK2^low^ populations. Column 2: Average NPAT or FLASH mRNA FISH puncta number and 95% confidence interval as a function of time since anaphase. (H) NPAT or FLASH mRNA FISH puncta number for CDK2^inc^ vs CDK2^low^ cells. p-values are indicated as four stars p ≤ 0.0001 and one star p ≤ 0.05.

Several key factors in histone biosynthesis, including Cyclin E, NPAT, FLASH, and SLBP, are transcriptional targets of cell-cycle master transcription factor E2F (Figure 1A) (Fischer et al., 2016; Gao et al., 2003; Sokolova et al., 2017). In quiescent cells, E2F is bound and inhibited by the retinoblastoma protein, Rb. Hyper-phosphorylation of Rb by cell-cycle master kinase CDK2/Cyclin E at the point of cell-cycle commitment known as the Restriction Point causes the release and activation of E2F (Figure 1A) (Hinds et al., 1992; Pardee, 1974; Zarkowska and Mittnacht, 1997). Cyclin E/CDK2 is also known to phosphorylate NPAT, with this phosphorylation being required for transcriptional activation of histone genes at the HLB (Figure 1A) (Gao *et al*., 2003; Hur et al., 2020; Ma et al., 2000; Zhao et al., 1998; Zhao et al., 2000). We previously showed that in unperturbed asynchronously cycling cells, a majority of cells will be born with elevated CDK2 activity having already crossed the Restriction Point (Moser *et al*., 2018; Spencer *et al*., 2013). We further demonstrated that in these cells born committed to the cell cycle, histone biosynthesis occurs in two phases, with low levels beginning at the Restriction Point and lasting throughout G1, followed by a burst of histone biosynthesis at the G1/S boundary (Armstrong and Spencer, 2021). However, it remains unclear if Cyclin E/CDK2 triggers the low level of histone transcription at the Restriction Point, if it triggers the burst at the start of S phase, or if the burst of histone production is triggered by stabilization of histone mRNA.

Once transcribed, histone mRNA is subject to secondary regulation via the degradation of mature histone mRNA through destabilization of the interaction between SLBP and the 3’ stem loop (Marzluff and Koreski, 2017; Sullivan *et al*., 2009). It has been previously shown that histone mRNA will be rapidly degraded in response to inhibition of DNA synthesis, in part due to the increase in soluble histone protein that has not been incorporated into chromatin (Kaygun and Marzluff, 2005a; b). Soluble histone protein must remain bound to histone chaperones from the time it is made until it is incorporated in the chromatin, or it will be subject to degradation (Hogan and Foltz, 2021; Jimeno-Gonzalez et al., 2015; Ransom et al., 2010). One such histone chaperone is nuclear autoantigenic sperm protein (NASP), which protects the available soluble histone H1, H3, and H4 pool from degradation (Bao et al., 2022; Cook et al., 2011; Finn et al., 2008). It remains unclear if soluble histone protein regulates histone mRNA stability in naturally cycling cells, or if this mechanism occurs only in response to DNA synthesis inhibition. Additionally, it is unknown if soluble histone protein’s ability to promote histone mRNA degradation is exclusive to S phase, or if the burst of histone mRNA at the G1/S boundary is due in part to excess soluble histones being incorporated into the chromatin as DNA replication begins.

In this study, we use single-cell timelapse imaging of asynchronously cycling non-transformed human MCF10A cells to determine the regulation of HLB formation, transcriptional activation of histone genes, and histone mRNA stabilization across the cell cycle. We find that HLB formation is initiated at the Restriction Point and is equally dependent on both NPAT and FLASH protein. Further, we find that phosphorylation of NPAT by Cyclin E/CDK2 at the Restriction Point triggers the initiation of histone transcription, which leads to the burst of histone mRNA at the G1/S boundary approximately 5 hr later. By contrast, phosphorylation of NPAT does not alter HLB formation. Finally, our data demonstrate that as cells enter S phase, regulation of histone mRNA levels transfers from CDK2-mediated transcriptional activation to degradation of histone mRNA via negative feedback from excess soluble histone protein. Therefore, cells reach an equilibrium between overlapping mechanisms to coordinate the burst in histone biosynthesis with the start of S phase, and then fine tune histone production in response to the rate of DNA replication.

## Results

### NPAT and FLASH are required for HLB formation

To map the temporal dynamics of HLB formation within the cell cycle, we tracked individual cells using single-cell timelapse imaging of asynchronously cycling cells, followed by post hoc staining of EdU and HLB factors, as has been previously described (Armstrong and Spencer, 2021; Gookin et al., 2017; Min and Spencer, 2019). We first transduced MCF10A cells with fluorescent histone 2B (H2B) as a nuclear marker, and a fluorescent fragment of DNA helicase B (DHB) as a sensor of CDK2 activity, measured by cytoplasmic to nuclear ratio of DHB (Figure 1B) (Spencer *et al*., 2013). Timelapse imaging, in which cells can be distinguished as proliferative (CDK2^inc^) or quiescent (CDK2^low^) based on DHB translocation, was followed by paraformaldehyde fixation of the cell, photobleaching of the DHB sensor, and staining for DNA content, EdU, NPAT as a marker for the HLB, and either FLASH or Lsm11 protein (Figure 1A-C). Cells were then aligned to the time of anaphase and classified as either CDK2^inc^ or CDK2^low^. CDK2^inc^ cells were further segmented by cell-cycle phase, as G1 (2N DNA content, EdU^neg^), S (EdU^pos^), or G2/M (4N DNA content, EdU^neg^) (STAR Methods) (see also Figure S1A-D).

HLB growth (as measured by the total size of all the HLBs per cell, or as the NPAT concentration within those HLBs) is initiated as early as 2 hr after anaphase in CDK2^inc^ cells and progresses continuously through G1 and S phase before plateauing in G2. (Figure 1C-D). We then quantified the concentration of FLASH or Lsm11 protein at the HLB, to map the recruitment of these factors relative to cell-cycle phase (Figure 1C-D). We found that FLASH is recruited in tandem with NPAT to the HLB, while Lsm11 recruitment remains lower throughout G1, before rising steadily in S phase and plateauing in G2/M (Figure 1D). This indicates that there may be a hierarchical recruitment of processing factors to the HLB as a means to regulate the timing of histone mRNA production, as has been previously suggested in *Drosophila* (Sabath et al., 2013; Skrajna *et al*., 2017).

To determine the dependency of HLB formation on each of these essential HLB proteins, we performed a 48 hr siRNA knockdown of NPAT, FLASH, and Lsm11, with live-cell imaging in the final 18 hr before fixation and HLB staining (Figure 1E; see also Figure S1E-F). We found that knockdown of either NPAT or FLASH significantly reduced HLB formation, with each protein being unable to localize to the HLB without the other. By contrast, knockdown of Lsm11 caused only a small reduction in HLB size and no reduction in protein concentrations of NPAT or FLASH, both of which increased slightly (Figure 1E). Additionally, we found that in cells that entered a CDK2^low^ quiescent state following mitosis, the HLB would reform initially and then dissipate over time (Figure S1G). Protein concentrations within the HLB in CDK2^low^ cells stayed fairly consistent over time for NPAT and FLASH, while Lsm11 concentration increased slightly as the HLBs got smaller, indicating that the HLB condensate retains HLB factors until it has fully dissipated (Figure S1G). Together, these data suggest that FLASH has an equally important role in seeding the HLB as NPAT, and that the HLB will form in cells that have both NPAT and FLASH protein available even if that cell is newly quiescent, lacks CDK2 activity, and is not producing histone mRNA.

Since NPAT and FLASH are both E2F target genes, we theorized that their expression would begin to rise immediately in CDK2^inc^ cells, since hyper-phosphorylation of Rb and the subsequent release of E2F is driven by CDK2 activity (Figure 1A). Following live-cell imaging, we stained for either NPAT or FLASH mRNA by RNA FISH along with a cytoplasmic marker and quantified the number of FISH puncta per cell (Figure 1F). We then aligned the single cells relative to their time of anaphase in CDK2^inc^ and CDK2^low^ cells (Figure 1G-H). We found that both NPAT and FLASH expression are significantly different between CDK2^inc^ and CDK2^low^ cells as early as 2 hr following mitosis, and that this difference in mRNA levels between CDK2^inc^ and CDK2^low^ cells is largely sustained throughout the cell cycle (Figure 1G-H). Thus, we conclude that HLB formation is fundamentally driven by E2F release at the Restriction Point, and the subsequent rise in NPAT and FLASH protein. Following crossing of the Restriction Point, HLB formation is dependent on NPAT and FLASH concentration.

### Lsm11 is recruited to the HLB after NPAT and FLASH have initiated condensate formation

To better quantify recruitment of Lsm11 and FLASH to the HLB, we performed high resolution 3D imaging on cells stained for DNA, NPAT, and either FLASH or Lsm11 (Figure 2A-B). Individual HLB puncta were segmented as 3D objects based on the NPAT signal, with subsequent FLASH and Lsm11 quantification and segmentation (STAR Methods). For individual HLBs, we quantified FLASH puncta volume relative to NPAT puncta volume and found a strong linear correlation between the size of NPAT puncta and FLASH puncta within the HLB, with FLASH puncta being consistently slightly smaller than NPAT puncta (Figure 2C). This may be due to the FLASH antibody recognizing the C-terminus of the protein, which has been shown to localize more strongly to the center of the HLB compared with the N-terminus of the protein (Kemp et al., 2021). Accurate volumetric measurements of Lsm11 HLB puncta were not possible due to the puncta being diffraction limited (STAR Methods).

**Figure 2.**
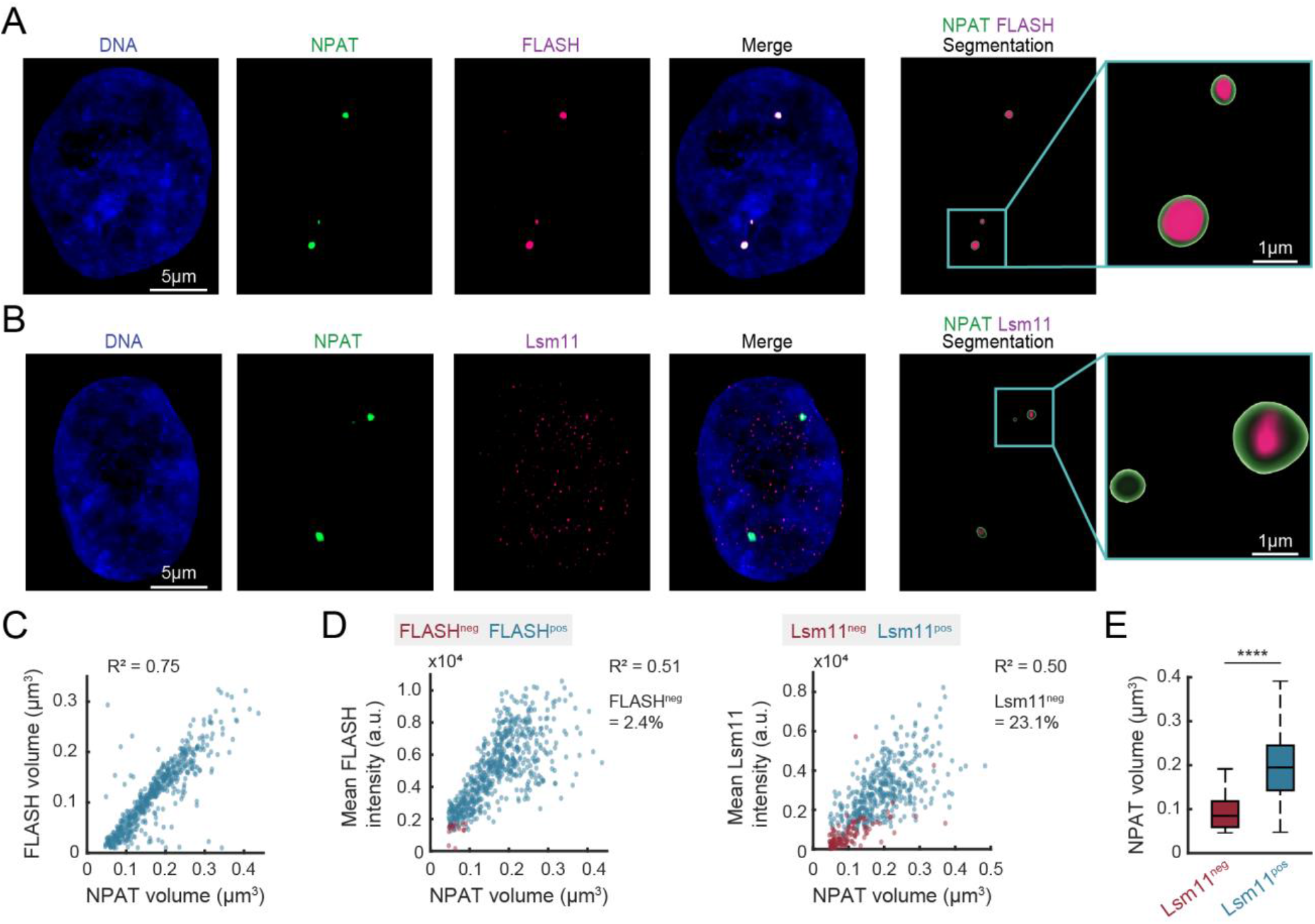
Lsm11 is recruited to the HLB after NPAT and FLASH have initiated condensate formation. (A-B) Representative images of cells stained for DNA, NPAT, and either FLASH (A) or Lsm11 (B). (C) Scatter of individual HLBs with FLASH volume vs NPAT volume. (D) Mean protein intensity for FLASH (column 1) or Lsm11 (column 2) vs NPAT volume of HLBs, designated as positive or negative for corresponding detection of either FLASH or Lsm11. (E) NPAT volume for HLBs that are Lsm11^neg^ or Lsm11^pos^. p-values are indicated as four stars p ≤ 0.0001.

Since the Lsm11 puncta were too small to accurately quantify their volume, we measured FLASH and Lsm11 recruitment to the HLB as mean protein intensity for FLASH or Lsm11 within the HLB, relative to NPAT puncta volume (Figure 2D). We further categorized HLB puncta as those with (FLASH^pos^/Lsm11^pos^) or without (FLASH^neg^/Lsm11^neg^) detection for FLASH or Lsm11 protein (Figure 2D) (STAR Methods). Localization of both FLASH and Lsm11 protein to the HLB correlates linearly with HLB growth, however, a significant percentage (23.1%) of the HLBs did not have corresponding Lsm11 signal (Figure 2D). Lack of Lsm11 occurred in the smallest HLBs, indicating that Lsm11 is recruited after HLB formation has been initiated by NPAT and FLASH (Figure 2E). It is unclear if Lsm11 localizes to the HLB once the condensate has reached a certain size, or if recruitment occurs later in the cell cycle after HLB growth has already been initiated.

### Transcriptional activation of histone genes at the HLB is triggered by Cyclin E/CDK2’s phosphorylation of NPAT

To gain further insight into the temporal dynamics and regulation of transcriptional activation of histone genes at the HLB, we tracked localization of Cyclin E to the HLB. We began by determining which forms of Cyclin E were accumulating in the HLB, by staining for both Cyclin E1 and Cyclin E2 using antibodies we validated by siRNA knockdown (Figure S2A-D), along with DNA, EdU, and NPAT (Figure 3A). We found that the pan-nuclear signal of both Cyclin E1 and E2 peaked approximately at the G1/S phase boundary as has been previously reported, with Cyclin E1 but not E2 continuing to rise in CDK2^low^ cells (Figure 3B) (since Cyclin E1 is not being actively degraded in the CDK2^low^ state) (Gookin *et al*., 2017; Min and Spencer, 2019). We then determined whether Cyclin E1 or E2 was specifically accumulating at the HLB by taking the average signal at the HLB as marked by NPAT and normalizing it to the nuclear signal for each protein (Figure 3C). We find that both Cyclin E1 and E2 have a pool of protein that localizes specifically to the HLB, with the highest levels in late-G1 and early-S, thereby enabling CDK2 to phosphorylate NPAT locally and in a cell-cycle dependent manner.

**Figure 3.**
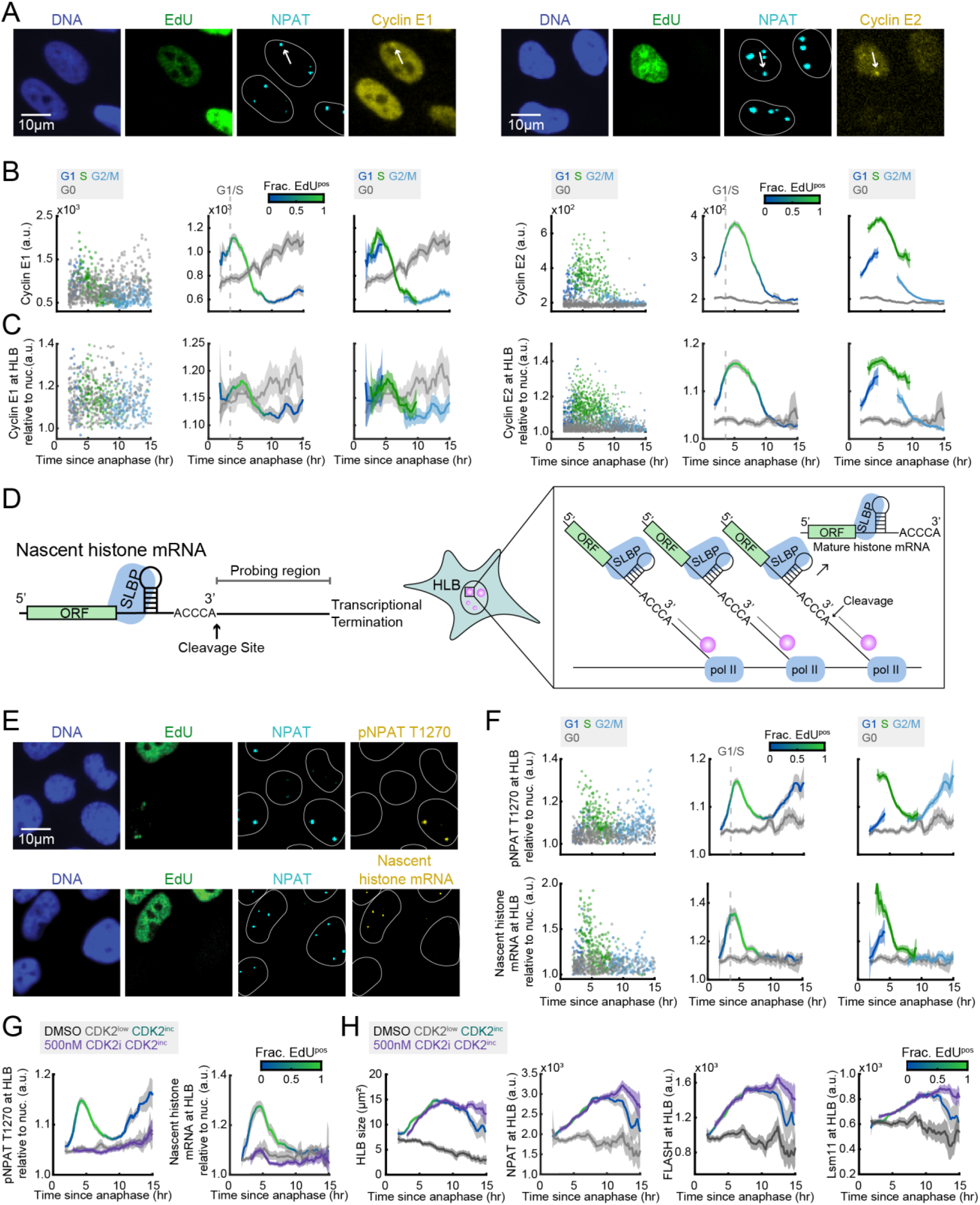
Transcriptional activation of histone genes at the HLB is triggered by Cyclin E/CDK2 phosphorylation of NPAT. (A) Representative images following timelapse imaging of cells stained for DNA, EdU, NPAT, and either Cyclin E1 or Cyclin E2. (B-C) Column 1: Raw single-cell data by cell-cycle phase. Column 2: Average nuclear signal for Cyclin E1 or Cyclin E2 (B) and average HLB signal for Cyclin E1 or Cyclin E2 normalized to nuclear signal (C) and 95% confidence intervals as a function of time since anaphase for CDK2^inc^ and CDK2^low^ populations with G1/S transition as time when 50% of cells are EdU^pos^. Column 3: CDK2^inc^ population from column 2 segmented in to G1, S, and G2/M. (D) Schematic of nascent histone mRNA FISH probe design (see Table S1). (E) Representative images following timelapse imaging of cells stained for DNA, EdU, NPAT, and either pNPAT T1270 or nascent histone mRNA FISH. (F) Column 1: Raw single-cell data by cell-cycle phase. Column 2: Average HLB signal for pNPAT T1270 or nascent histone mRNA normalized to nuclear signal and 95% confidence intervals as a function of time since anaphase for CDK2^inc^ and CDK2^low^ populations with G1/S transition as time when 50% of cells are EdU^pos^. Column 3: CDK2^inc^ population from column 2 segmented in to G1, S, and G2/M. (G) Average HLB signal for pNPAT T1270 and nascent histone mRNA FISH normalized to nuclear signal for CDK2^inc^ and CDK2^low^ populations for cells treated with DMSO, and CDK2^inc^ population for cells treated with 500nM PF-07104091 (CDK2i) for the last 1 hr of live-cell imaging. (H) Average HLB size and average HLB signal with 95% confidence interval for NPAT, FLASH, and Lsm11 for CDK2^inc^ and CDK2^low^ populations for cells treated with DMSO, and CDK2^inc^ population for cells treated with 500nM PF-07104091 (CDK2i) in the for the last 1 hr of live-cell imaging.

To look more directly at transcriptional activation at the HLB, we developed a custom antibody against phospho-NPAT (pNPAT) T1270 based on a previously published epitope (Ma *et al*., 2000). We also designed a probe set against nascent histone mRNA to quantify transcriptional activation of histone genes at the HLB via RNA FISH (STAR Methods, see also Table S1). Since histones genes are intron-less, we designed probes against the sequence downstream of the 3’ cleavage site and prior to transcriptional termination for 50 of the histone genes found at the HIST1 and HIST2 clusters (Figure 3D, see also Table S1) (Marzluff *et al*., 2002). Cells were stained for DNA, EdU, NPAT, and either pNPAT T1270 or nascent histone RNA FISH following timelapse imaging (Figure 3E). We normalized pNPAT T1270 and nascent histone signals to the nuclear background in cells that have HLBs since both are HLB-specific signals (Figure 3F). Both pNPAT T1270 and nascent histone mRNA levels begin rising upon cell-cycle commitment, peak at the G1/S boundary, and fall to baseline levels at the end of S phase (Figure 3F). This coincides exactly with the timing of Cyclin E1’s rise and fall, as opposed to Cyclin E2, which falls several hours later (Figure 3B-C). Unexpectedly, there is then a second rise in pNPAT T1270 at the HLB that does not correlate with histone gene transcription, beginning at G2 and ending at mitosis. The role of this later wave of NPAT phosphorylation is unknown (Figure 3F). The strong correlation in timing of Cyclin E1, pNPAT T1270, and nascent histone mRNA suggests that activation and cessation of transcription at the HLB is predominantly regulated by Cyclin E1/CDK2 activity.

To further test this hypothesis, we utilized PF-07104091 (CDK2i), a novel and selective small molecule inhibitor of CDK2 (Fassl et al., 2022; Jhaveri et al., 2021; Mezi et al., 2021; Rana et al., 2021). Cells were treated with 500nM CDK2i during the last hour of live-cell imaging before fixing at staining for DNA, EdU, NPAT, and either pNPAT T1270 or nascent histone RNA FISH (Figure 3G, see also Figure S3A-B). CDK2 inhibition was verified by the nuclear translocation of the DHB sensor upon drug treatment (Figure S4A-B). We found that inhibition of Cyclin E/CDK2 led to an immediate and complete loss of both NPAT phosphorylation and nascent histone transcription (Figure 3G). To determine if this was due to the loss of HLB formation, we treated cells with 500nM CDK2i during the last hour of live-cell imaging and stained for DNA, EdU, NPAT, and either FLASH or Lsm11 (Figure 3H, see also Figure S3C-D). HLB formation was unchanged with CDK2i treatment, indicating that formation of the HLB and initiation of histone transcription are decoupled once the cells have crossed the Restriction Point. These effects were consistent at lower and higher doses of CDK2i (100nM and 2µM) (Figure S3A-F). Thus, we find that histone transcription is positively regulated by Cyclin E/CDK2 phosphorylation of NPAT from the Restriction Point to the S/G2 transition, peaking at the G1/S boundary.

### Histone mRNA degrades rapidly and proportionally upon inhibition of DNA synthesis

Having shown that transcription of histone genes is triggered by Cyclin E/CDK2 activity upon cell-cycle commitment and that the rate of mature histone mRNA production peaks at the G1/S phase boundary, we next sought to understand if there was a secondary regulation mechanism suppressing excess histone biosynthesis in a cell-cycle dependent manner. We began by establishing the expression dynamics of the histone mRNA stabilizing protein, SLBP, and histone H4.2 mRNA relative to the cell cycle (Figure 4A-B, see also Figure S2E and H). The results were consistent with our previously published findings that SLBP levels are high throughout G1 and S phase and fall at the S/G2 boundary (Figure 4C) (Armstrong and Spencer, 2021). Quantification of SLBP mRNA similarly shows elevated expression in CDK2^inc^ vs CDK2^low^ cells (Figure S5A-C). Additionally, histone mRNA is sustained at low-to-moderate levels throughout G1 before a rapid accumulation of histone mRNA at the G1/S phase boundary, coincident with EdU incorporation, as we and others showed previously (Figure 4D) (Armstrong and Spencer, 2021; Harris et al., 1991; Heintz et al., 1983). These data indicate that in addition to SLBP abundance, histone mRNA stabilization must depend on an alternative mechanism more closely linking histone production with DNA synthesis in S phase.

**Figure 4.**
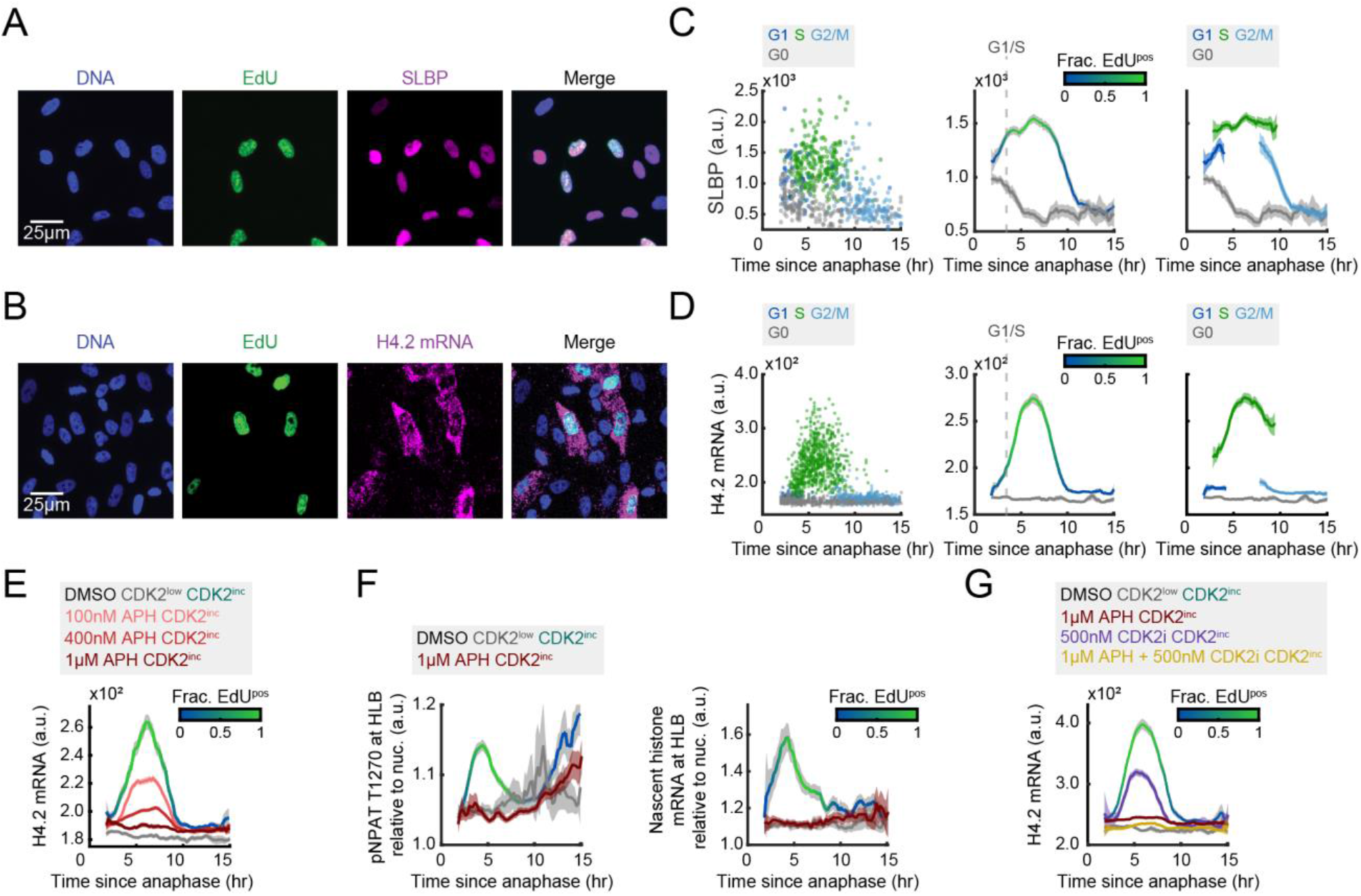
Histone mRNA degrades rapidly and proportionally upon inhibition of DNA synthesis. (A-B) Representative images following timelapse imaging of cells stained for DNA, EdU, and SLBP (A) or H4.2 mRNA (B). (C-D) Column 1: Raw single-cell data by cell-cycle phase. Column 2: Average nuclear SLBP signal (C) or cytoplasmic histone H4.2 mRNA (D) and 95% confidence intervals as a function of time since anaphase for CDK2^inc^ and CDK2^low^ populations, with G1/S transition as the time when 50% of cells are EdU^pos^. Column 3: CDK2^inc^ population from column 2 segmented in to G1, S, and G2/M. (E) Average histone H4.2 mRNA signal for CDK2^inc^ and CDK2^low^ populations for cells treated with DMSO, and CDK2^inc^ population for cells treated with 100nM, 400nM, or 1µM APH for the last 1 hr of live-cell imaging. (F) Average HLB signal for pNPAT T1270 and nascent histone mRNA FISH normalized to nuclear signal for CDK2^inc^ and CDK2^low^ populations for cells treated with DMSO, and CDK2^inc^ population for cells treated with 1µM APH for the last 1 hr of live-cell imaging. (G) Average histone H4.2 mRNA signal and 95% confidence interval for CDK2^inc^ and CDK2^low^ populations for cells treated with DMSO, and CDK2^inc^ population for cells treated with 1µM APH, 500nM PF-07104091 (CDK2i), or a combination of 1µM APH and 500nM PF-07104091 (CDK2i) for the last 1 hr of live-cell imaging.

To test this hypothesis, we inhibited DNA synthesis by treating cells with aphidicolin (APH) at a range of doses to inhibit DNA synthesis partially or completely (100nM, 400nM, and 1µM), for 1 hr before the end of live-cell imaging. We then fixed and stained the cells for DNA, EdU, and histone H4.2 mRNA (Figure 4E, see also Figure S5D-F). Histone H4.2 mRNA levels fell proportionally to the degree of DNA synthesis inhibition, regardless of the timing within S phase (Figure 4E), suggesting that cells have a mechanism by which they can immediately sense and respond to changes in the rate of DNA replication. To address if this change in histone mRNA was due to a loss of transcription or degradation of the mRNA, we treated cells with 1µM APH for 1 hr before staining for pNPAT T1270 or nascent histone mRNA FISH, and found that inhibition of DNA synthesis prevents new histone transcription (Figure 4F, see also Figure S5G-H). However, when we treated cells with 500nM CDK2i, which fully inhibits histone transcription (Figure 3G), histone mRNA is only partially reduced, suggesting that loss of histone mRNA with 1µM APH treatment is due at least in part to histone mRNA degradation (Figure 4G, see also Figure S5I-J). Reduction in histone mRNA upon 500nM CDK2i treatment is likely due to a decrease in DNA synthesis, as CDK2 activity is also needed for DNA replication to proceed (Figure S5I-J). We also noted that when treated cells with a combination of 1µM APH and 500nM CDK2i, histone mRNA levels fell even further than with 1µM APH treatment alone, despite DNA synthesis being fully inhibited in both conditions (Figure 4G, see also Figure S5I-J). This suggests that CDK2 activity might not only activate transcription of histones but may potentially also play a smaller role in protecting histone mRNA from degradation. By contrast, inhibition of DNA synthesis has no effect on HLB formation (Figure S6A-C) or SLBP levels (Figure S6D-E), demonstrating that the reduction histone mRNA is specifically due to loss of transcription and an increase in histone mRNA degradation. Together, these data indicate that cells respond to changes in DNA synthesis rate in S phase both by preventing further histone gene expression and actively degrading the histone mRNA that is already present.

### Soluble histone protein promotes the degradation of histone mRNA in S phase

Previous studies have suggested that soluble histone protein may promote the degradation of histone mRNA, but it is not known if this negative feedback regulates the burst of histone mRNA at the G1/S boundary (Figure 5A) (Kaygun and Marzluff, 2005b). To answer this question, we used siRNA to deplete cells of Eri1 or NASP. Knockdown of Eri1, the gene for histone mRNA-specific exonuclease 3’hExo, should prevent histone mRNA degradation, and result in an increase in histone mRNA. Partial knockdown of histone chaperone NASP provides a means to modulate the soluble histone pool of histones H1, H3, and H4 (Bao *et al*., 2022; Cook *et al*., 2011). NASP is histone chaperone that is not required for nucleosome remodeling within the chromatin, but instead specifically shields the soluble histone pool from degradation (Figure 5A) (Cook *et al*., 2011; Hormazabal et al., 2022). Therefore, knockdown of NASP should decrease the amount of soluble histone protein and increase the amount of histone mRNA.

**Figure 5.**
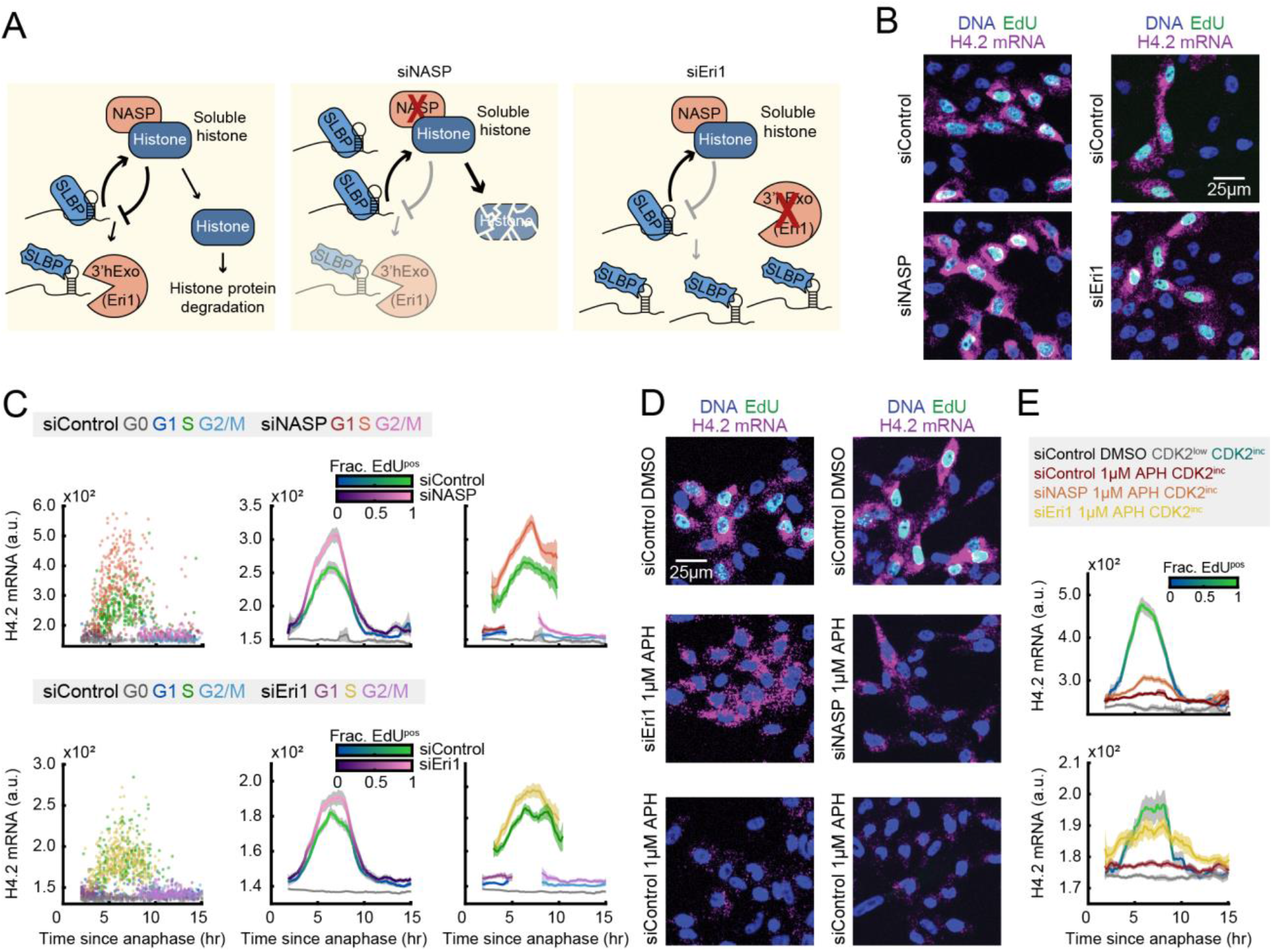
Soluble histone protein promotes the degradation of histone mRNA in S phase. (A) Schematic of histone protein and histone mRNA stabilization and siRNA knockdown of NASP and Eri1. (B) Representative images of cells treated with 20nM siControl or siNASP for 24 hr before fixation, or cells treated with 25nM siControl or siEri1 for 48 hr before fixation. (C) Column 1: Raw single-cell data. Column 2: Average cytoplasmic histone H4.2 mRNA signal and 95% confidence intervals as a function of time since anaphase for CDK2^inc^ and CDK2^low^ populations for cells treated with siControl vs siNASP or siControl vs siEri1. Column 3: CDK2^inc^ population from column 2 segmented in to G1, S, and G2/M populations. (D) Representative images of cells treated with 20nM siControl for 24 hr with 1 hr DMSO, 20nM siControl for 24 hr with 1 hr 1µM APH, or 20nM siNASP for 24 hr with 1 hr 1µM APH before fixation; and cells treated with 25nM siControl for 48 hr with 1 hr DMSO, 25nM siControl for 48 hr with 1 hr 1µM APH, or 25nM siEri1 for 48 hr with 1 hr 1µM APH before fixation. (E) Average histone H4.2 mRNA signal and 95% confidence interval for cells in (D).

To directly test if soluble histone protein can promote the degradation of histone mRNA, we partially knocked down NASP via siRNA for 24 hr, filming cells for the last 18 hr of the knockdown period, followed by fixing and staining for DNA, EdU, and histone H4.2 mRNA (Figure 5B). Similarly, we knocked down Eri1, via siRNA for 48hr prior to fixation, with the last 18 hr of the siRNA knockdown being filmed, followed by staining for DNA, EdU, and histone H4.2 mRNA (Figure 5B). The siRNA knockdown of both NASP and Eri1 were validated by immunofluorescent staining of NASP and 3’hExo (Figure S2F-G and I-J). We found that reduction of the soluble histone pool via partial knockdown of NASP led to higher levels of histone H4.2 mRNA in S phase specifically. Additionally, prevention of histone mRNA degradation via knockdown of Eri1 led to an increase in histone mRNA levels in S phase and a slight increase in histone mRNA levels in G1 (Figure 5C). Therefore, we find that soluble histone protein promotes histone mRNA degradation specifically in S phase, while an alternative mechanism promotes histone mRNA degradation in G1 to a lesser degree.

To further test whether the decrease of histone mRNA upon inhibition of DNA synthesis is caused by an increase in soluble histone protein, we paired the siRNA knockdown of NASP and Eri1 with a 1 hr treatment of 1µM APH prior to fixing and staining for DNA, EdU, and histone H4.2 mRNA (Figure 5D). In the presence of APH, Histone H4.2 mRNA levels were increased in S phase cells by partial knockdown of NASP and subsequent decrease in the soluble histone pool (Figure 5D-E). Additionally, histone H4.2 mRNA levels in the presence of APH were largely rescued via knockdown of Eri1, indicating that the loss in histone mRNA upon inhibition of DNA synthesis is predominantly due to degradation (Figure 5D-E). Rescue of histone mRNA following the knock down of NASP or Eri1 suggests that upon inhibition of DNA synthesis, soluble histone protein directly acts to promote the degradation of histone mRNA. Thus, as cells enter S phase, the incorporation rate of soluble histone into the chromatin can promote the degradation of histone mRNA, thereby balancing histone production with the rate of DNA synthesis.

## Discussion

In this study, we investigated the temporal dynamics of the mechanisms which regulate histone biosynthesis relative to the G1/S boundary. It was long thought that crossing of the Restriction Point and the start of S phase happened close in time, and thus that histone biosynthesis was tied solely with DNA replication (Marzluff et al., 2008; Zhao *et al*., 1998). Our recent studies, however, have illustrated that crossing of the Restriction Point occurs about 5 hr before the start of S phase, and thus these events are decoupled temporally (Spencer *et al*., 2013). Therefore, the mechanistic basis of the burst of histone biosynthesis at the G1/S boundary was not properly understood. Here, we demonstrate that histone biosynthesis undergoes positive and negative regulation from the Restriction Point to the start of S phase via three cell-cycle-dependent mechanisms: 1) formation of the HLB driven by E2F-mediated expression of NPAT and FLASH; 2) initiation of histone transcription upon phosphorylation of NPAT by Cyclin E/CDK2; and 3) the rate of nucleosome incorporation into chromatin during DNA replication dictating soluble histone protein levels, and therefore the rate of histone mRNA degradation in S phase. As cells transition from G1 to S phase, they reach an equilibrium between Cyclin E/CDK2-mediated transcriptional activation of histone genes and soluble histone protein-mediated degradation of histone mRNA, with both processes responding rapidly to loss of DNA replication. Therefore, the burst of mature histone mRNA observed at the start of S phase is caused by the initiation of nascent histone transcription by Cyclin E/CDK2’s phosphorylation of NPAT several hours earlier at the Restriction Point. The peak of nascent histone pre-mRNA at the G1/S boundary correlates temporally with the fastest rate of increase of mature histone mRNA, since the actively transcribed histone pre-mRNA must be processed before accumulating as mature histone mRNA, whereas the nascent pre-mRNA is transitory (Figure 6A).

**Figure 6.**
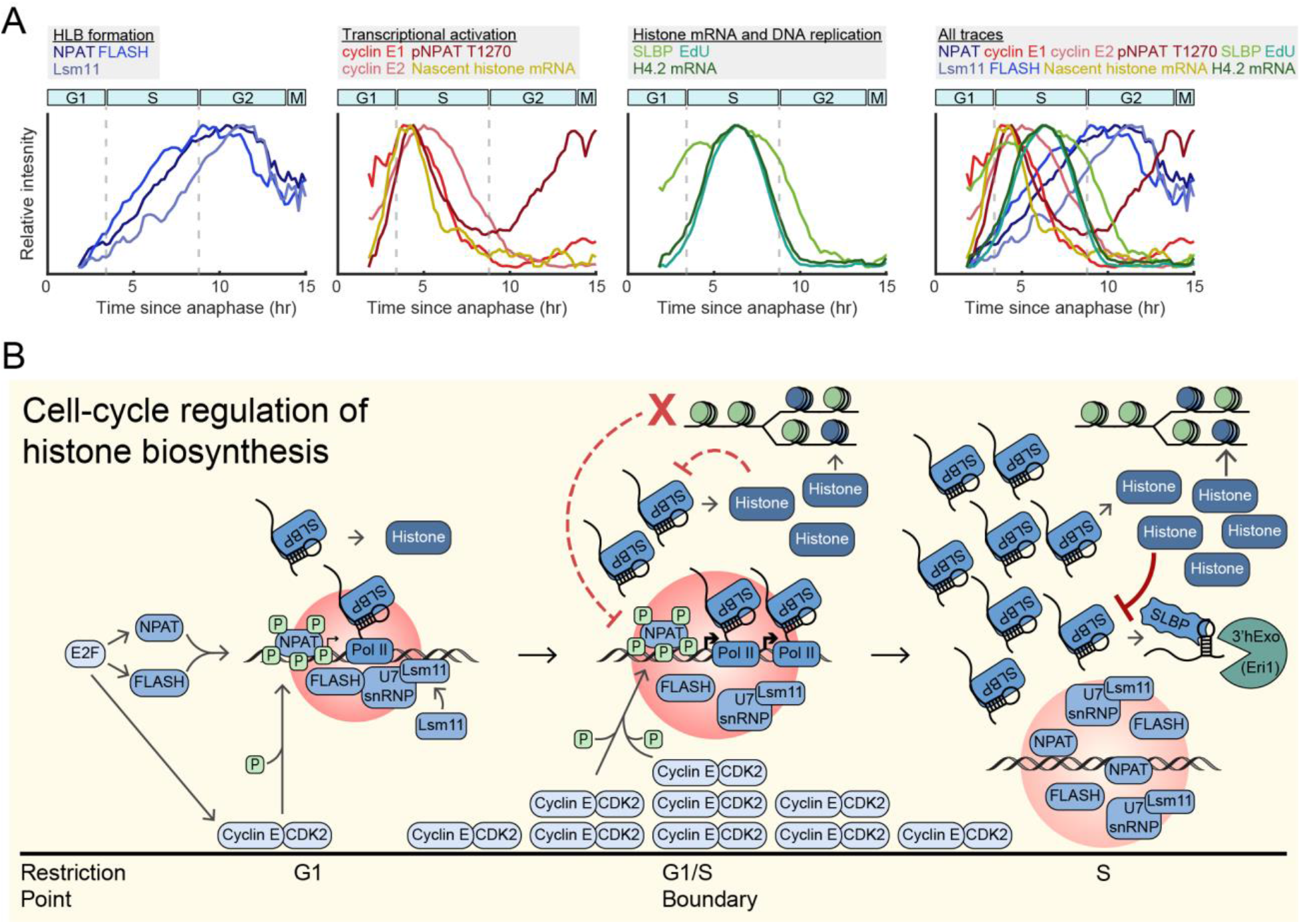
Temporal dynamics and cell-cycle regulation of histone biosynthesis. (A) Overlay of normalized average signals presented in this study, categorized by HLB formation, transcriptional activation, histone mRNA and DNA replication, and all traces. (B) Schematic of cell-cycle regulation of histone biosynthesis relative to cell-cycle progression.

### Decoupling of HLB Formation and Function

Formation of transcriptional condensates, such as super enhancers, myc condensates, and HLBs, are essential for the high levels of transcription and processing of the genes they envelope. However, the correlation between formation and activation of these transcriptional condensates has remained unclear. Here, we demonstrate that on short time scales, the phosphorylation of the HLB seeding and scaffolding protein NPAT activates transcription at the HLBs, but does not affect HLB formation, in contrast to what has been seen in Drosophila embryos (Hur *et al*., 2020). We further show that the cessation of transcription upon CDK2 inhibition does not occur through a dissociation of the HLB, but through loss of NPAT phosphorylation. Additionally, NPAT appears to undergo a later wave of phosphorylation in G2 whose functional role is unknown. Loss of this second phosphorylation event upon inhibition of CDK2 suggests that NPAT may be phosphorylated by Cyclin A/CDK2 specifically in G2. This late phosphorylation of NPAT late may play a role in HLB dissociation in mitosis, or in preparing the cell for histone biosynthesis in its subsequent cell cycle. Together, these data suggest that once cells have crossed the Restriction Point, HLB formation and transcription activation are decoupled in human cells, a mechanism which ensures prolonged stability of the HLB while tightly regulating the timing of histone gene transcription.

### Self-regulation of histone protein

Once histone mRNA has been transcribed and processed, it is no longer under the regulation of HLB. Yet high levels of histone production in the event of a slowing of DNA synthesis would be toxic to the cell due to the accumulation of excess soluble histone and the aberrant interactions they would form with negatively charged molecules (Hogan and Foltz, 2021). We find that cells address this via a negative-feedback mechanism in which excess soluble histone protein promotes the of degradation histone mRNA. This likely occurs through destabilization of the SLBP interaction with the 3’ stem loop, which normally blocks degradation of histone mRNA by 3’hExo (Marzluff and Koreski, 2017), although the precise mechanism of this destabilization is unknown. Activation of ATR upon inhibition of DNA synthesis has been found to destabilize the SLBP-3’ stem loop interaction via Upf1, as well as inhibit CDK2, likely causing the loss of histone transcription (Daigh et al., 2018; Kaygun and Marzluff, 2005a; Marzluff and Koreski, 2017; Meaux *et al*., 2018). Additionally, it was surprising to find that soluble histone protein suppresses histone mRNA even when DNA synthesis is unperturbed. These data indicate that this is a mechanism in naturally cycling cells to keep histone biosynthesis in step with DNA synthesis (Figure 6A). Our work maps the timing by which various mechanisms regulate histone biosynthesis as cells transition from the point of cell-cycle commitment to the end of DNA replication (Figure 6B), thereby precisely detailing one of the most critical processes for faithful replication of the genome.

### Limitations of the study

While we were able to use single-cell live-cell imaging linked with immunofluorescent and RNA FISH staining to measure each protein and RNA of interest at each time point relative to each cell’s last mitosis, a limitation of this study is absence of live-cell sensors for our histone biosynthesis factors. Development of a live-cell sensor for NPAT, as exists for its *Drosophila* homolog Mxc, would allow for tracking of HLB growth and formation in live single cells (Hur *et al*., 2020; Terzo et al., 2015). Observing HLB growth in individual cells would provide deeper understanding of the heterogeneity of HLB formation that may exist within the population. Quantifying HLB formation in tandem with other live-cell sensors for other HLB factors, SLBP, or expression of histone genes, would serve to validate the temporal dynamics presented herein and determine if variation in histone biosynthesis correlates with changes in DNA replication stress or cell-cycle fate.

Additionally, due to the challenging and laborious nature of these experiments, the results presented in this study were performed only in MCF10A cells. To test the universality of these dynamics, the histone biosynthesis would ideally be mapped in several cell types, including normal primary cells and cancer cells. This would provide a comparison between healthy and aberrant histone biosynthesis dynamics.

## Acknowledgements

We thank Francesca Mattiroli for insights on the project and feedback on the manuscript, as well as the members of the laboratory of S.L.S. for general help and feedback on the manuscript. This work was supported by NIH training grant T32 (GM065103-16) (to C.A.), Pew-Stewart Scholar for Cancer Research Award, an American Cancer Society Research Scholar Grant (RSG-18-008-01), and an NIH Director’s New Innovator Award (1DP2CA238330-01) (to S.L.S).

## Author contributions

Conceptualization C.A.; Methodology C.A. and V.J.P.; Validation C.A. and V.J.P.; Formal Analysis C.A., H.M.A., and V.J.P.; Investigation C.A. and V.J.P.; Resources C.A., V.J.P, S.L.S.; Data Curation C.A. and V.J.P.; Writing-Original Draft C.A. and V.J.P.; Writing-Review & Editing C.A., V.J.P., S.L.S.; Visualization C.A.; Supervision S.L.S.; Project Administration S.L.S.; Funding Acquisition S.L.S.

## STAR Methods

### Method Details

#### Cell Culture and Maintenance

MCF10A were obtained from ATCC (CRL-10317) and were cultured in Dulbecco’s Modified Eagle Medium/Nutrient Mixture F-12 (DMEM/F12) supplemented with 5% horse serum, 100 ng/mL cholera toxin, 20 ng/mL EGF, 10 µg/mL insulin, 0.5 µg/mL hydrocortisone, and 100 µg/mL of both penicillin and streptomycin. All cells were cultures at 37°C with 5% CO_2_. Live-cell imaging of MCF10A cells was done using a phenol-red free version of the growth media (Thermo Fisher, 11039047).

#### Antibodies

Primary antibodies were used at the following dilutions: NPAT (sc-136007 1:400, 611344 1:4000), SLBP (ab221166, 1:1000), FLASH (HPA053573, 1:4000), Lsm11 (PA5-31324, 1:500), Eri1 (14592-1-AP, 1:250), NASP (HPA030520, 1:2000), Cyclin E1 (ab33911, 1:4000), Cyclin E2 (ab40890, 1:500), pNPAT T1270 (1:2000). Secondary antibodies were used at a dilution of 1:500, with the exception of FLASH, which was stained with a secondary dilution of 1:1000.

#### Custom antibody development

The antibody against phospho-NPAT T1270 is a polyclonal rabbit antibody produced by the Thermo Fisher custom antibody services. The construct was designed as previously described, with the sequence Asp-Leu-Pro-Val-Pro-Arg-phosphoThr-Pro-Gly-Ser-Gly-Ala-Gly-Cys, generated after coupling to keyhole limpet hemocyanin (Ma *et al*., 2000).

#### Immunofluorescent staining

Cells were fixed with 4% paraformaldehyde for 15 minutes, washed 3 times with PBS, then permeabilized in 0.1% TritonX for 20 minutes. Standard protocols were then used for immunofluorescent staining: cells were first blocked in 3% BSA for 1 hour at 37°C, primary antibodies were incubated in 3% BSA overnight at 4°C, cells were washed three times in PBS, secondary antibodies were then incubated at room temperature for 2 hours, cells were washed three times in PBS before being incubated with Hoechst at 1:10,000 at room temperature for 20 minutes. Imaging was done on a Nikon Ti-E with a 10x 0.45 NA objective with the appropriate filter applied. Exposure times were set to 400ms for DAPI, 300ms for Cy3, and 400ms for Cy5.

#### RNA FISH staining

Cells were fixed with 4% paraformaldehyde for 15 minutes. Histone H4.2 (Thermo Fisher, VA6-3174283-VC), NPAT (Thermo Fisher, VA6-3172751-VC), FLASH (Thermo Fisher, VA6-3175253-VC), and SLBP (Thermo Fisher, VA6-3174137-VCP) mRNA were visualized according to the manufacturer’s protocol (ViewRNA ISH Cell Assay Kit, ThermoFisher QVC0001), with cells being permeabilized for 30 minutes and mRNA probes hybridized for 4 hours at 40°C. Exposure times were set to 600ms for Cy5.

#### Nascent histone RNA FISH design and staining

##### Probe design

We designed probes targeted against the sequence downstream of the 3’ cleavage site for 50 histone genes found at clusters HIST1 and HIST2 (see Table S1 for sequence and position relative to the cleavage site), to measure nascent histone mRNA expression at the HLB. Probes were designed with the Stellaris RNA FISH Probe Designer (Biosearch Technologies, Inc., Petaluma, CA) available online at www.biosearchtech.com/stellarisdesigner (Version 4.2). Probes were set to have an oligo length of 20 nt, a masking level of 5, and a minimum spacing length of 2nt. Starting positions were chosen to be within 25nt of the cleavage site for HIST1, and 150nt of the cleavage site for HIST2.

##### Probe set labeling

Probes for nascent histone RNA FISH were ordered from IDT. Each oligo was resuspended at a concentration of 250µM in nuclease free H_2_O, before being pooled in an oligo master mix. Probes were then labeled with ATTO 550 using ddUTP-ATTO550 (Jena Bioscience, NU-1619-550) with terminal deoxynucleotidyl transferase (Thermo Fisher, EP0161), as previously described (Gaspar et al., 2017).

##### Staining

Cells were fixed with 4% paraformaldehyde for 15 minutes, before being permeabilized with 0.1% TritonX for 30 minutes. Nascent histone RNA was then visualized in accordance with the manufacture’s protocol (https://biosearch-cdn.azureedge.net/assetsv6/protocol_stellaris-adherent-cells-96-well-glass-bottom-plates.pdf) (Biosearch Technologies, Inc., Petaluma, CA; Wash Buffer A, SMF-WA1-60; Wash Buffer B, SMF-WB1-20; Hybridization Buffer, SMF-HB1-10). Labeled nascent histone mRNA probes were hybridized for 16 hr at 37°C at a dilution of 1:50. Exposure times were set to 800ms for Cy3. Following nascent histone RNA FISH staining and imaging, cells were stained for NPAT protein as described above.

#### siRNA transfection

siRNA transfections were performed using the DharmaFECT 1 (Dharmacon, #T-2001-02) reagent as described by the manufacturer. For live-cell imaging, the transfection mix was added to cells with 20nM final concentration for siNASP and 25nM final concentration for siEri1, siNPAT, siFLASH, and siLsm11, and removed after 6 hr. Imaging was started immediately and proceeded for an additional 18 hr for siNASP, or imaging began 24 hr after transfection mixture was removed and proceeded for an additional 18 hr for siEri1, siNPAT, siFLASH, and siLsm11. For the validation of antibodies for Cyclin E1, Cyclin E2, and SLBP, transfection mixture was added to cells with 25nM final concentration of siRNA, was removed after 6 hr, with cells being fixed and stained as described above after 24 hr for siCCNE1 and siCCNE2, and after 48hr for siSLBP. Oligonucleotides used in this study include: control (Horizon, D-001810-02-05), NPAT (Horizon, J-019599-10-0002), SLBP (Horizon, J-012286-05-0002), FLASH (Horizon, J-012413-05-0002), Lsm11 (Horizon, J-018455-17-0002), Eri1 (Horizon, J-021497-09-0002), NASP (Horizon, J-011740-13-0002), CCNE1 (Horizon, M-003213-02-0005), and CCNE2 (Horizon, M-003214-02-0005).

#### Live-cell imaging

##### Live-cell movies with HLB imaging

Cells were plated on a 96-well plate (Cellvis Cat. No. P96-1.5H-N) coated with collagen at a 1:50 dilution in water (Advanced BioMatrix, #5005). Cells were plated at densities of 3,600 cells per well 24 hr prior to the start of imaging for all movies expect for those with 48 hr siRNA knockdowns, which were plated at 1,600 cells per well 48 hr prior to the start of imaging. All live cell-imaging was done on a Nikon Ti-E microscope with a 10x 0.45 NA objective, except for movies followed by staining for nascent histone mRNA, which were done with a 20x 0.45 NA objective, with the appropriate filter applied, at a frequency of 5 frames per hour. For the duration of the movie, cells were maintained in a humidified incubation chamber at 37°C, with 5% CO2. Exposure times were set to 70ms per frame for CFP, corresponding to H2B-mTurqiose2, and 100ms for mCherry, corresponding to DHB-mCherry. Prior to the last frame of imaging, all cells were pulsed with 10µM of EdU for 12 minutes when applicable, before being immediately fixed with 4% paraformaldehyde. Following permeabilization with 0.1% TritonX for 20 minutes, cells were photobleached with 3% H_2_O_2_ for 20 minutes until the DHB-mCherry signal was no longer present. Cells were then processed for EdU visualization as described by the manufacturer’s protocol (Invitrogen, C10340) with Alexa fluor 488. Subsequent immunofluorescent or nascent histone mRNA FISH staining was done as described above. Exposure times were set to 400ms for DAPI, 600ms for YFP, 300ms for Cy3, and 400ms for Cy5 unless otherwise specified.

##### All other live-cell movies

Cells were plated as described above 24 hr prior to the start of imaging at a density of 3,600 cells per well for all movies expect for those with 48 hr siRNA knockdowns, which were plated at 1,600 cells per well 48 hr prior to the start of imaging. Live cell-imaging was done on a Nikon Ti-E microscope with a 10x 0.45 NA objective with the appropriate filter applied, at a frequency of 5 frames per hour. For the duration of the movie, cells were maintained in a humidified incubation chamber at 37°C with 5% CO2. Exposure times were set to 70ms per frame for CFP, corresponding to H2B-mTurqiose2, and 100ms for YFP, corresponding to DHB-mVenus. Prior to the last frame of imaging cell were pulsed with 10µM of EdU for 12 minutes when applicable, before being immediately fixed with 4% paraformaldehyde, and processed for EdU visualization as described by the manufacturer’s protocol (Invitrogen, C10340) with Alexa fluor 555. Cells were then processed for RNA FISH or immunofluorescent staining, as described above. Exposure times were set to 400ms for DAPI, 200ms for Cy3, and 400ms for Cy5, unless otherwise specified.

### 3D imaging

Following immunofluorescent staining, cells were imaged using a Nikon A1R Laser Scanning Confocal microscope equipped with an Andor iXon 897 Ultra EMCCD and NIS Elements software (v. 5.30). Cells with detectable HLB puncta were selected using a 10x NA 0.45 air objective and their XY coordinates were saved. Individual Z-stacks of selected cells were then captured sequentially using a Plan Apo λ 100x 1.45 NA oil objective with 0.1 µm step size and 1.0 AU pinhole size per channel. Chromatic aberration between the TRITC and Cy5 channels was quantified using 0.5 µm fluorescent microspheres on a FocalCheck test slide (Invitrogen, F36909). The Cy5 channel was computationally shifted by +0.06 µm in the X plane for all nuclear Z-stacks to compensate for the measured aberration.

### Quantification and Statistical Analysis Tracking of live-cell imaging

Image processing, cell-tracking, and cell classification were done as previously described (Arora et al., 2017; Gookin *et al*., 2017; Spencer *et al*., 2013), with the tracking code available at https://github.com/scappell/Cell_tracking. Cyclin E1, Cyclin E2, and SLBP protein signals, as well as EdU incorporation, were quantified as the median pixel value of the nuclear mask, histone H4.2 mRNA FISH signal was quantified as the median pixel value of a four-pixel-wide cytoplasmic ring around the nuclear mask. Protein or mRNA signals were matched back to the last frame of the live-cell movie using nearest neighbor screening after jitter correction. HLB puncta were identified from NPAT images, as previously described (Armstrong and Spencer, 2021; Arora *et al*., 2017). The MATLAB function regionprops was used label all puncta and to retrieve the xy coordinates of all pixels belonging to all the puncta of a given nucleus. Pre-imaging of the nascent histone mRNA FISH was linked to post-imaging of NPAT and EdU in the same cells by matching of nuclear centroids. Average protein or RNA FISH signal at the HLB was determined as the sum of the protein signal of all puncta pixels per cell normalized to the HLB size, with the HLB size being defined as the total number of puncta pixels per cell. For signals where nuclear protein content did not inherently determine localization to the HLB, Cyclin E1, Cyclin E2, pNPAT T1270, and nascent histone mRNA FISH, the average signal at the HLB in cells with HLB puncta was normalized to the median nuclear intensity.

### Classification of cell populations

Following alignment to anaphase, cells were classified as either CDK2^inc^ or CDK2^low^ based on the level of CDK2 activity 2 hr after anaphase. Cells with CDK2 activity above 0.5 were classified as CDK2^inc^, while cells that stayed under 0.5 for the remainder of imaging were classified as CDK2^low^, followed by removing any EdU^pos^ or 4N DNA content cells from the population. CDK2^inc^ cells were further classified by cell-cycle phase, with G1 cells as 2N DNA content, EdU^neg^; S cells as EdU^pos^; G2/M cells as 4N DNA content, EdU^neg^ (see also Figure S1A-D). CDK2^inc^ vs CDK2^low^ classification was extended to 3 hr after anaphase for cells treated with PF-07104091 (CDK2i), and the duration of time required for cells to have a CDK2 activity above 0.5 to be classified as CDK2^inc^ was shorted from 4 hr to 0.2 hr, since CDK2i treatment caused the DHB sensor to translocate back to the nucleus from the cytoplasm (see Figure S4A-B).

### HLB quantification from 3D imaging

Laser scanning confocal deconvolution was applied to each Z-stack using the Richardson-Lucy deconvolution algorithm for 20 iterations using Imaris software (v. 9.9.0). Nuclei were segmented for 3D object creation using a local contrast threshold and filtered with a minimum diameter gate of 15 µm. NPAT and FLASH puncta within the nucleus were segmented using a local contrast threshold and filtered for volumes above the diffraction limit of the imaging system. Lsm11 foci were detected using the Imaris Spots function as they were too small for accurate volume measurement. The distances between centroids of all segmented NPAT and FLASH or NPAT and Lsm11 puncta were calculated and colocalization events were called when the calculated distance was at or below the resolution limit of the imaging system.

## Data and Software Availability

Raw live-cell movies and corresponding images for Figures 1-6 were deposited on Mendeley at DOI: 10.17632/d2b55pmjmk.1.

