## Supplemental info for "Cyclin E/CDK2 and feedback from soluble histone protein regulate the S phase burst of histone biosynthesis"

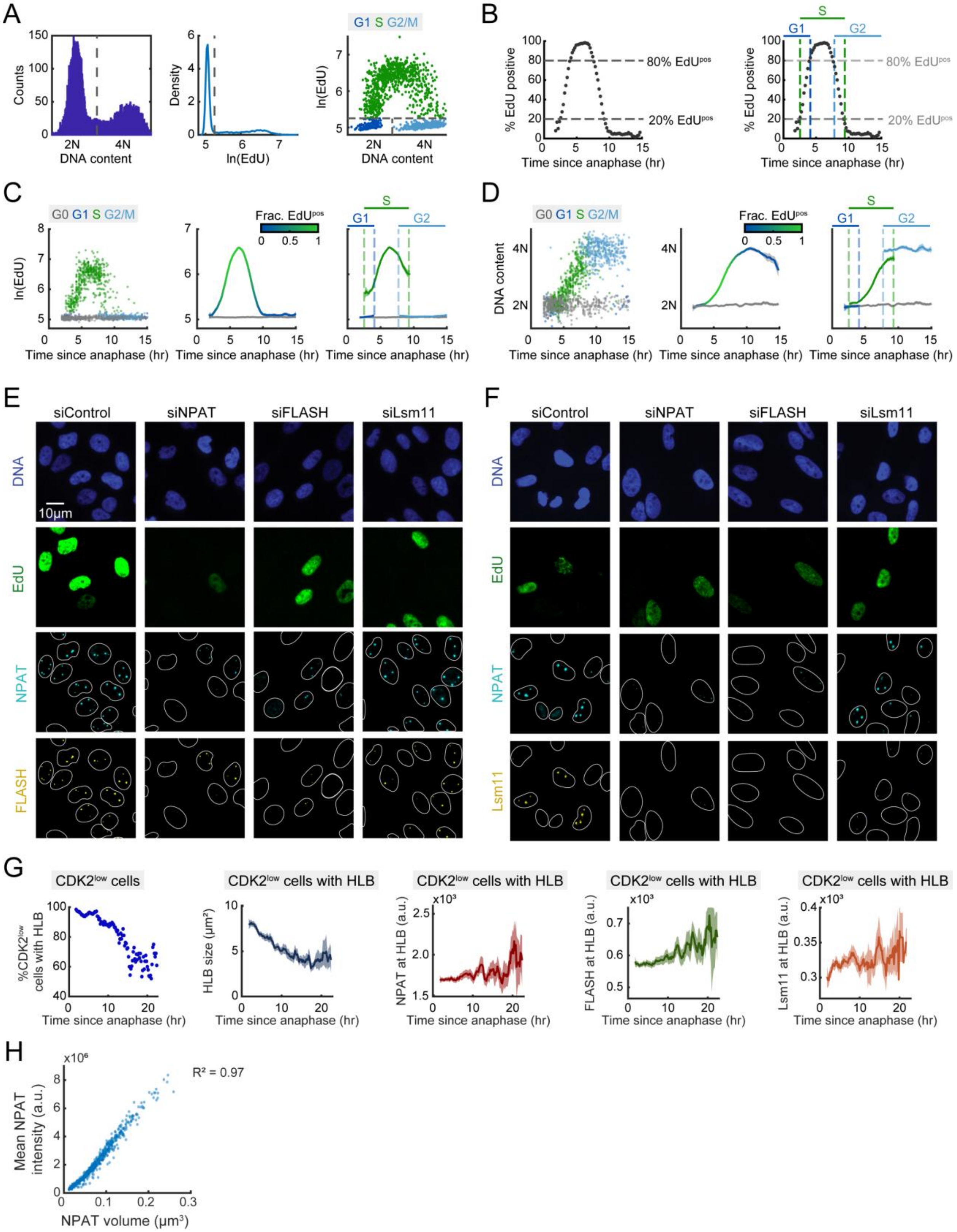


**Figure S1. Determination of cell-cycle phase, representative images of cells treated with siControl, siNPAT, siFLASH, and siLsm11, and HLB quantification in CDK2$^{low}$ cells, related to Figure 1-2.** (A)

Column 1: Histogram of DNA content based on Hoestch signal of all cells in the population, with cutoff for 2N vs 4N DNA content being determined by local minima. Column 2: ksdensity plot of EdU signal for all cells in the population, with cutoffs for $EdU^{neg}$ vs. $EdU^{pos}$ being determined by local minima. Column 3: Representative scatter of EdU vs. DNA content in $CDK2^{inc}$ cells following timelapse imaging segmented by cell-cycle phase. (B) Column 1: Percent $EdU^{pos}$ cells at each timepoint for $CDK2^{inc}$ population aligned to anaphase. Column 2: Cutoff of timing for each cell-cycle phase, where G1 cells are plotted from anaphase to the timepoint when 80% of cells are $EdU^{pos}$; S cells from the timepoint when 20% of cells are $EdU^{pos}$ until the timepoint when 80% of cells are $EdU^{neg}$; and G2 cells between the timepoint when 20% of cells are $EdU^{neg}$ until the end of the cell cycle. (C-D) Column 1: Raw single-cell data. Column 2: Average EdU (C) or DNA content (D) signal and 95% confidence intervals as a function of time since anaphase for $CDK2^{inc}$ and $CDK2^{low}$ populations. Column 3: $CDK2^{inc}$ population from column 2 segmented into G1 cells defined as 2N DNA content and $EdU^{neg}$, S cells defined as $EdU^{pos}$, and G2/M populations defined as 4N DNA content and $EdU^{neg}$. (E-F) Representative images of cells treated with 25nM of siControl, siNPAT, siFLASH, and siLsm11 for 48 hr prior to fixation with live-cell imaging in the final 18 hr, stained for DNA, EdU, NPAT, and either FLASH (E) or Lsm11 (F) (see Figure 1E). (G) Column 1: Percent of $CDK2^{low}$ cells with HLB puncta as a function of time spent out of the cell cycle. Column 2-5: Average HLB size (2) and average HLB signal for NPAT (3), FLASH (4), or Lsm11 (5) with 95% confidence interval in $CDK2^{low}$ cells with HLBs. (H) Scatter of mean NPAT intensity vs NPAT volume.

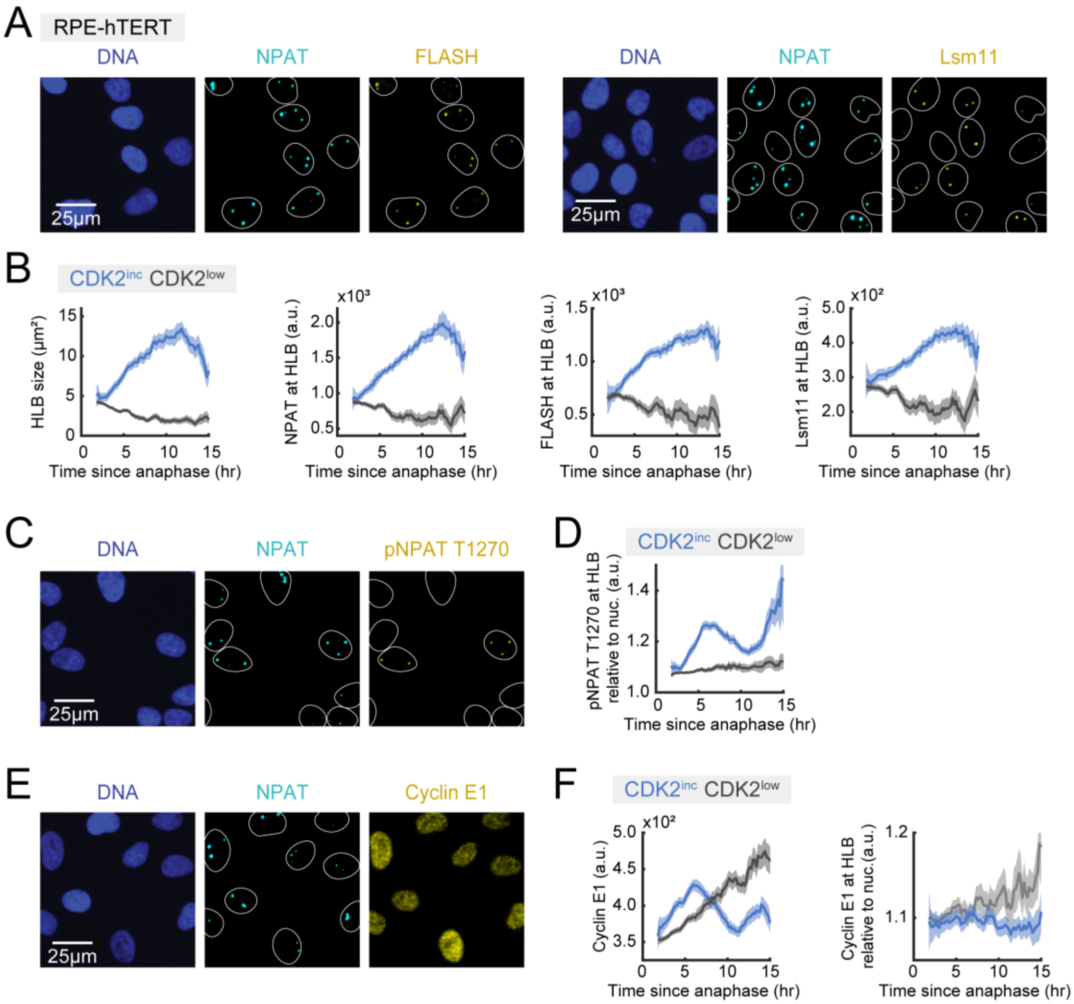


**Figure S2. HLB formation and NPAT phosphorylation in RPE-hTERT cells related to Figure 1-3.** (A) Representative images of cells following timelapse imaging, stained for DNA content, NPAT and either FLASH or Lsm11. (B) Average of the total HLB size per cell or protein signal at the HLB and 95% confidence intervals as a function of time since anaphase for CDK2$^{inc}$ and CDK2$^{low}$ populations. (C) Representative images of cells following timelapse imaging, stained for DNA content, NPAT and pNPAT T1270. (D) Average HLB signal for pNPAT T1270 normalized to nuclear signal for CDK2$^{inc}$ and CDK2$^{low}$ populations. (E) Representative images of cells following timelapse imaging, stained for DNA content, NPAT and Cyclin E1. (F) Average nuclear signal for Cyclin E1 (left) and average HLB signal for Cyclin E1 normalized to nuclear signal (right) and 95% confidence intervals as a function of time since anaphase for CDK2$^{inc}$ and CDK2$^{low}$ populations.

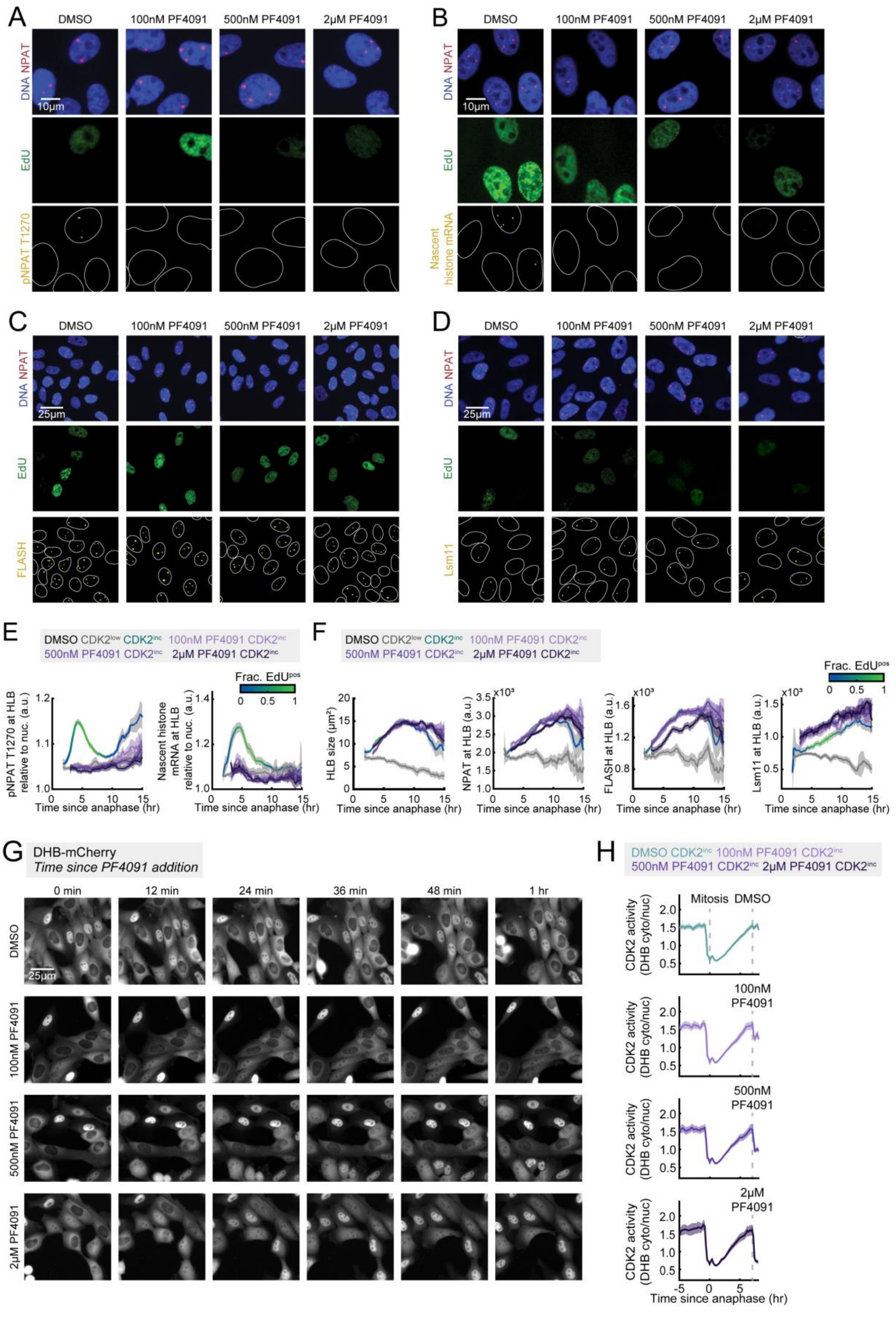

A
DMSO
100nM PF4091
500nM PF4091
2μM PF4091
DNA NPAT
10μm
EdU
pNPAT T1270
B
DMSO
100nM PF4091
500nM PF4091
2μM PF4091
DNA NPAT
10μm
EdU
Nascent histone mRNA
C
DMSO
100nM PF4091
500nM PF4091
2μM PF4091
DNA NPAT
25μm
EdU
FLASH
D
DMSO
100nM PF4091
500nM PF4091
2μM PF4091
DNA NPAT
25μm
EdU
Lsm11
E
DMSO CDK2low CDK2inc 100nM PF4091 CDK2inc
500nM PF4091 CDK2inc 2μM PF4091 CDK2inc
Frac. EdUpos
pNPAT T1270 at HLB relative to nuc. (a.u.)
Nascent histone mRNA at HLB relative to nuc. (a.u.)
Time since anaphase (hr)
F
DMSO CDK2low CDK2inc 100nM PF4091 CDK2inc
500nM PF4091 CDK2inc 2μM PF4091 CDK2inc
Frac. EdUpos
HLB size (μm²)
NPAT at HLB (a.u.)
FLASH at HLB (a.u.)
Lsm11 at HLB (a.u.)
Time since anaphase (hr)
G
DHB-mCherry
Time since PF4091 addition
0 min
12 min
24 min
36 min
48 min
1 hr
DMSO
100nM PF4091
500nM PF4091
2μM PF4091
25μm
H
DMSO CDK2inc 100nM PF4091 CDK2inc
500nM PF4091 CDK2inc 2μM PF4091 CDK2inc
Mitosis
DMSO
100nM PF4091
500nM PF4091
2μM PF4091
CDK2 activity (DHB cyto/nuc)
Time since anaphase (hr)

**Figure S3. Representative images and additional data on PF4091 treatment, related to Figure 3.** (A-D) Representative images of cells treated with 100nM, 500nM, and 2µM of PF-07104091 (PF4091) for the last 1 hr of live-cell imaging, before staining for DNA, EdU, NPAT, and either pNPAT T1270 (A), nascent histone mRNA (B), FLASH (C) or Lsm11 (D). (E) Average HLB signal for pNPAT T1270 and nascent histone mRNA FISH normalized to nuclear signal for $CDK2^{inc}$ and $CDK2^{low}$ populations for cells treated with DMSO, and $CDK2^{inc}$ population for cells treated with 100nM, 500nM, or 2µM PF-07104091 (PF4091) for the last 1 hr of live-cell imaging. (F) Average HLB size and average HLB signal with 95% confidence interval for NPAT, FLASH, and Lsm11 for $CDK2^{inc}$ and $CDK2^{low}$ populations for cells treated with DMSO, and $CDK2^{inc}$ population for cells treated with 100nM, 500nM, or 2µM PF-07104091 (PF4091) in the for the last 1 hr of live-cell imaging. (G) Representative images of the final 1 hr of live-cell imaging of the CDK2 activity sensor in cells treated with 100nM, 500nM, or 2µM PF-07104091 (PF4091). (H) Average CDK2 activity and 95% confidence interval of $CDK2^{inc}$ cells aligned to anaphase that completed their last mitosis 10 hr before the final frame of the movie, with drug added in the final 1 hr of imaging (see Figure 3D, 3F).

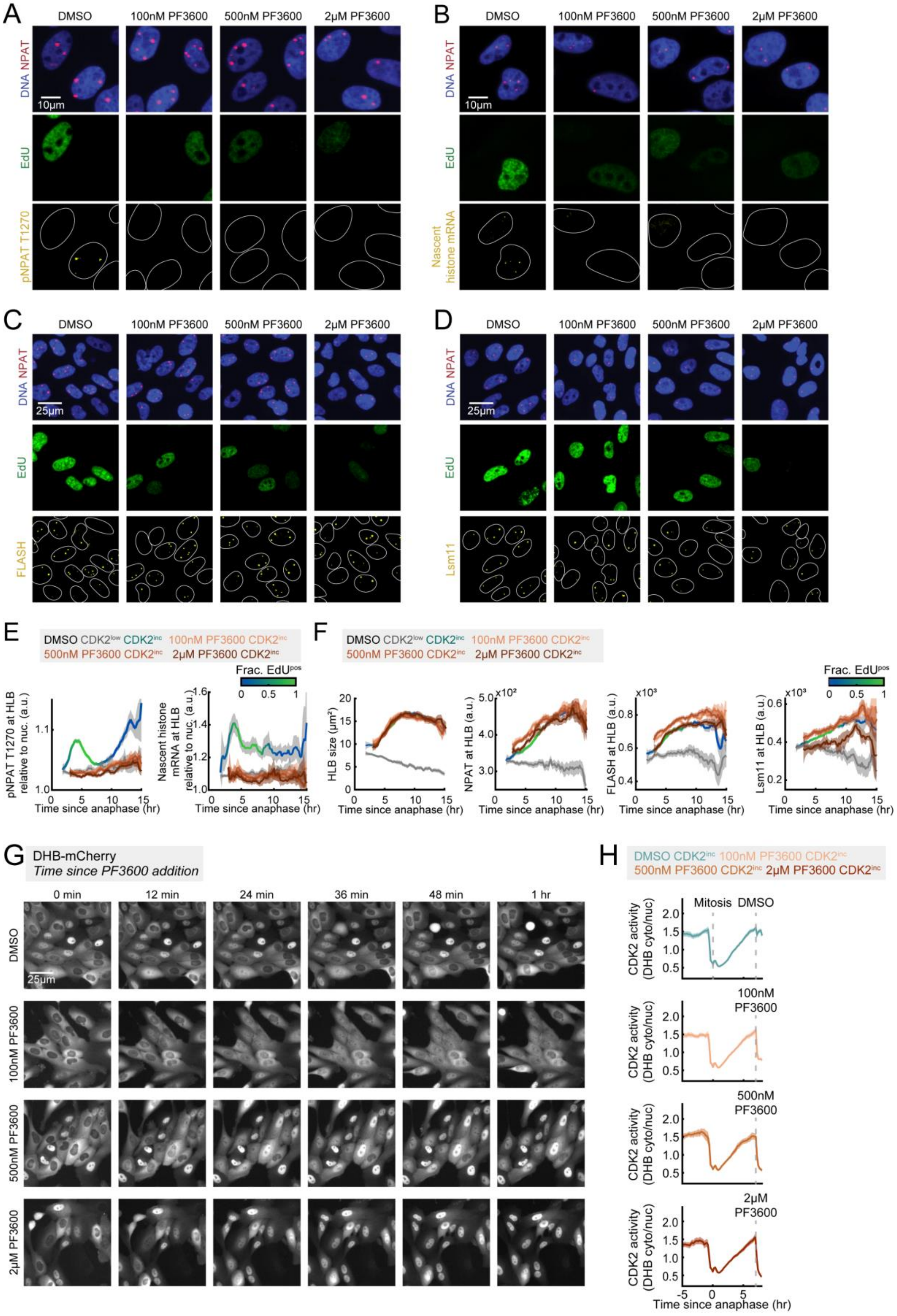
A
DMSO
100nM PF3600
500nM PF3600
2μM PF3600
DNA NPAT
10μm
EdU
pNPAT T1270
B
DMSO
100nM PF3600
500nM PF3600
2μM PF3600
DNA NPAT
10μm
EdU
Nascent histone mRNA
C
DMSO
100nM PF3600
500nM PF3600
2μM PF3600
DNA NPAT
25μm
EdU
FLASH
D
DMSO
100nM PF3600
500nM PF3600
2μM PF3600
DNA NPAT
25μm
EdU
Lsm11
E
DMSO CDK2low CDK2inc 100nM PF3600 CDK2inc
500nM PF3600 CDK2inc 2μM PF3600 CDK2inc
Frac. EdUpos
pNPAT T1270 at HLB relative to nuc. (a.u.)
Nascent histone mRNA at HLB relative to nuc. (a.u.)
Time since anaphase (hr)
F
DMSO CDK2low CDK2inc 100nM PF3600 CDK2inc
500nM PF3600 CDK2inc 2μM PF3600 CDK2inc
Frac. EdUpos
HLB size (μm²)
NPAT at HLB (a.u.)
FLASH at HLB (a.u.)
Lsm11 at HLB (a.u.)
Time since anaphase (hr)
G
DHB-mCherry
Time since PF3600 addition
0 min
12 min
24 min
36 min
48 min
1 hr
DMSO
25μm
100nM PF3600
500nM PF3600
2μM PF3600
H
DMSO CDK2inc 100nM PF3600 CDK2inc
500nM PF3600 CDK2inc 2μM PF3600 CDK2inc
Mitosis
DMSO
100nM PF3600
500nM PF3600
2μM PF3600
CDK2 activity (DHB cyto/nuc)
Time since anaphase (hr)

**Figure S4. DHB sensor translocation during PF3600 treatment, related to Figure 3.** (A-D) Representative images of cells treated with 100nM, 500nM, and 2µM of PF-06873600 (PF3600) for the last 1 hr of live-cell imaging, before staining for DNA, EdU, NPAT, and either pNPAT T1270 (A), nascent histone mRNA (B), FLASH (C) or Lsm11 (D). (E) Average HLB signal for pNPAT T1270 and nascent histone mRNA FISH normalized to nuclear signal for $CDK2^{inc}$ and $CDK2^{low}$ populations for cells treated with DMSO, and $CDK2^{inc}$ population for cells treated with 100nM, 500nM, or 2µM PF-06873600 (PF3600) for the last 1 hr of live-cell imaging. (F) Average HLB size and average HLB signal with 95% confidence interval for NPAT, FLASH, and Lsm11 for $CDK2^{inc}$ and $CDK2^{low}$ populations for cells treated with DMSO, and $CDK2^{inc}$ population for cells treated with 100nM, 500nM, or 2µM PF-06873600 (PF3600) in the for the last 1 hr of live-cell imaging. (G) Representative images of the final 1 hr of live-cell imaging of the CDK2 activity sensor in cells treated with 100nM, 500nM, or 2µM PF-06873600 (PF3600). (H) Average CDK2 activity and 95% confidence interval of $CDK2^{inc}$ cells aligned to anaphase that completed their last mitosis 10 hr before the final frame of the movie, with drug added in the final 1 hr of imaging (see Figure 3E, 3G).

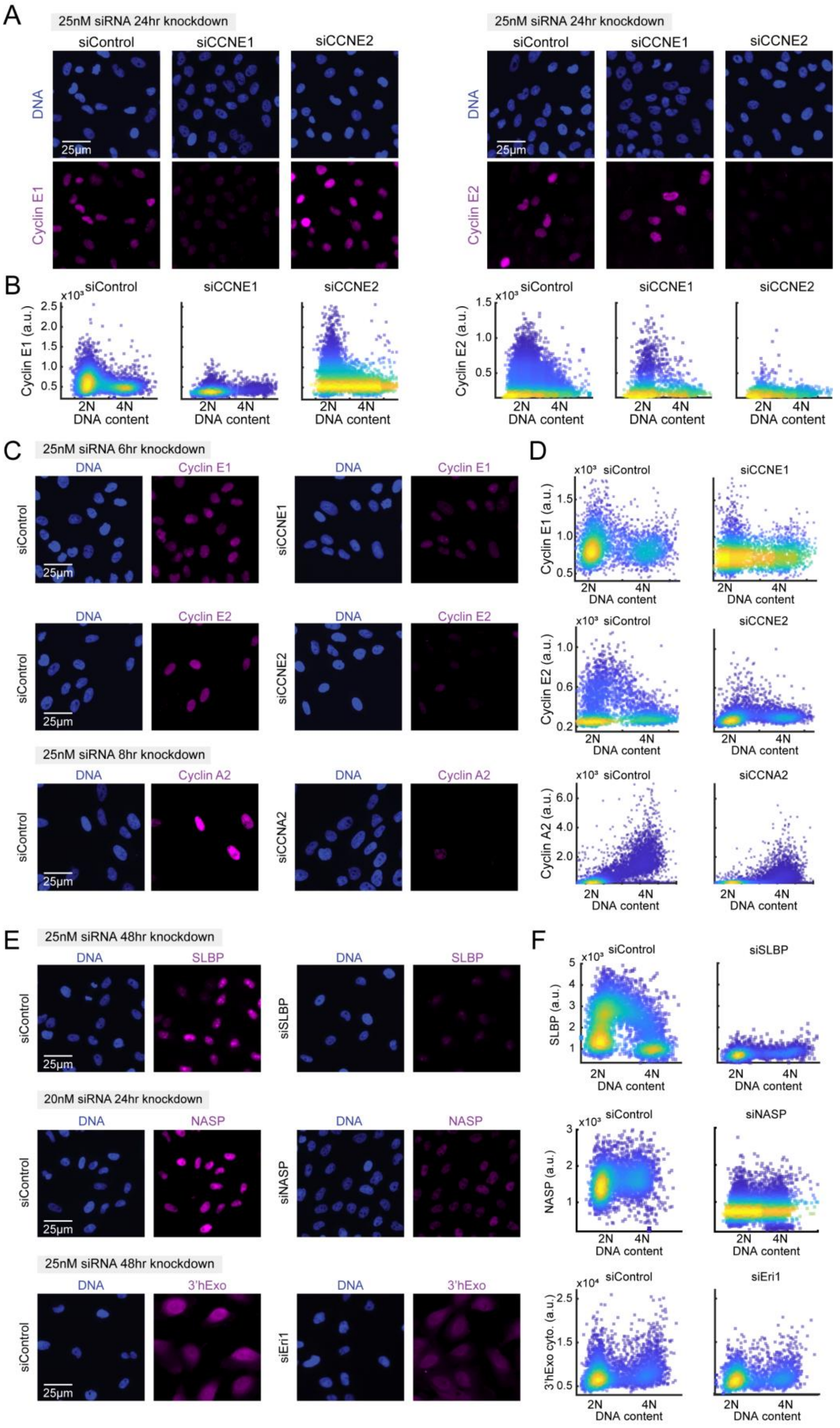
A
25nM siRNA 24hr knockdown
siControl
siCCNE1
siCCNE2
DNA
Cyclin E1
Cyclin E2
25µm
B
Cyclin E1 (a.u.)
Cyclin E2 (a.u.)
DNA content
2N
4N
C
25nM siRNA 6hr knockdown
25nM siRNA 8hr knockdown
Cyclin A2
siCCNA2
D
Cyclin A2 (a.u.)
E
25nM siRNA 48hr knockdown
20nM siRNA 24hr knockdown
SLBP
siSLBP
NASP
siNASP
3'hExo
siEri1
F
SLBP (a.u.)
NASP (a.u.)
3'hExo cyto. (a.u.)

**Figure S5. Validation of antibodies and siRNA knockdowns, related to Figure 4-6.** (A-B) Staining for DNA and either Cyclin E1 (A) or Cyclin E2 (B) in cells treated with siControl, siCCNE1, or siCCNE2 (see Figure 4A-C). (C-D) Representative images (C) and single cells plotted as Cyclin E1, Cyclin E2, or Cyclin A2 vs. DNA content (D) for cells treated with siControl, siCCNE1, or siCCNE2 for 6 hr, or siControl or siCCNA2 for 8 hr (see Figure 4E). (E-F) Representative images (E) and single cells plotted as SLBP, NASP, or 3'hExo vs. DNA content (F) in cells treated with siControl and either siSLBP, siNASP, or siEri1 (see Figure 5A, 6B-G).

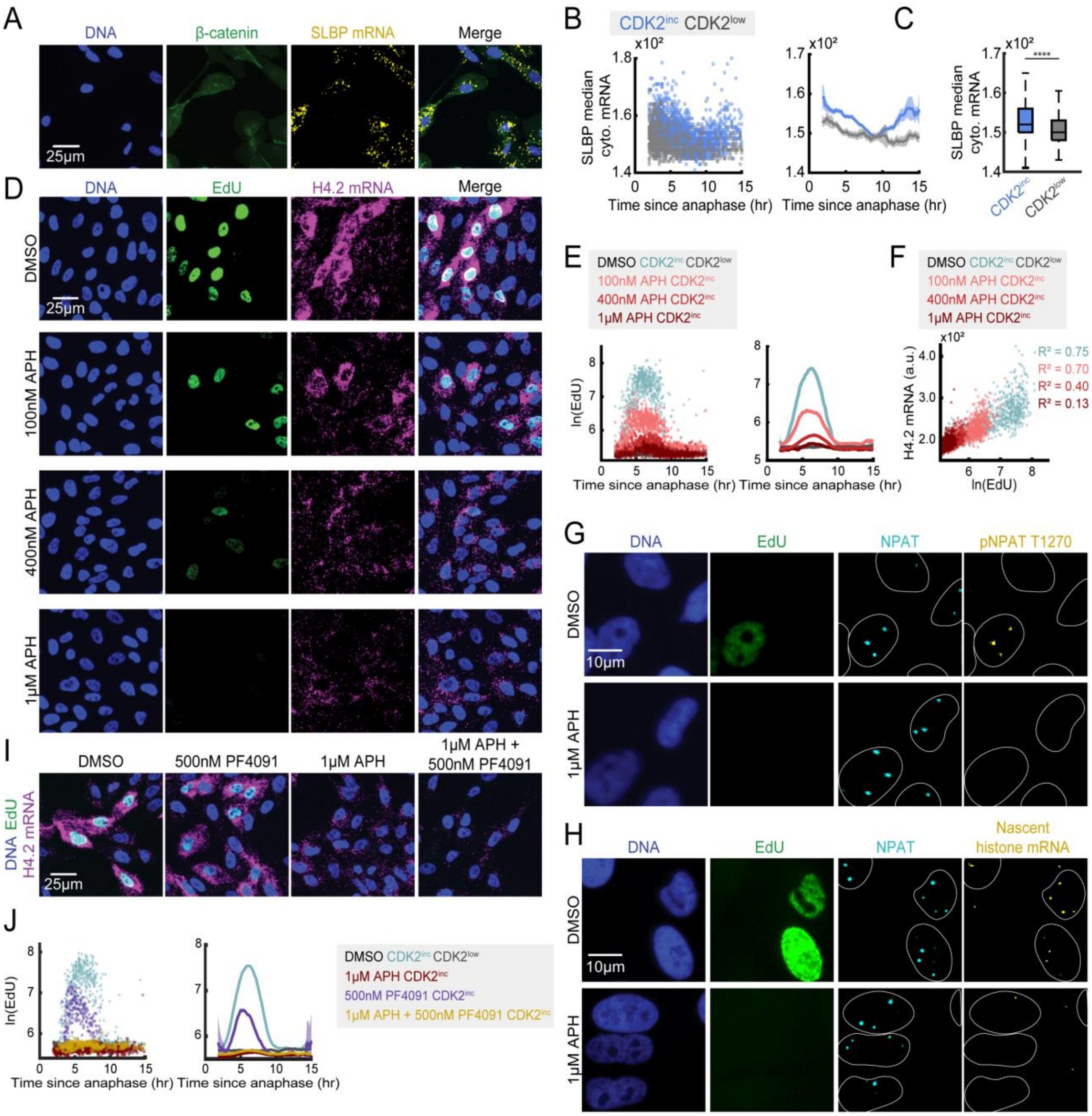


**Figure S6. SLBP expression and EdU incorporation upon DNA synthesis and CDK2 inhibition, related to Figure 5.** (A) Representative images of cells following live cell imaging, stained for DNA, β-catenin as a cytoplasmic marker, and SLBP mRNA. (B) Column 1: Raw single-cell data for $CDK2^{inc}$ and $CDK2^{low}$ populations. Column 2: Average cytoplasmic signal for SLBP mRNA and 95% confidence interval as a function of time since anaphase for data in column 1. (C) Median for $CDK2^{inc}$ vs $CDK2^{low}$ cells of SLBP mRNA. p-values are indicated as four stars $p \leq 0.0001$. (D) Representative images of cells treated with DMSO, 100nM, 400nM, or 1µM APH during the last 1 hr of live-cell imaging, before staining for DNA, EdU, and histone H4.2 mRNA. (E) Column 1: Raw single-cell data. Column 2: Average EdU signal and 95% confidence intervals as a function of time since anaphase for $CDK2^{inc}$ and $CDK2^{low}$ cells treated with DMSO, as well as $CDK2^{inc}$ cells treated with 100nM, 400nM, or 1µM APH (see Figure 5E). (F) Representative scatter of histone H4.2 mRNA vs. EdU of $CDK2^{inc}$ cells for cells treated with DMSO, 100nM APH, 400nM APH, or 1µM APH, with the R-squared value reported. (G-H) Representative images of cells treated with DMSO or 1µM APH in the last 1 hr of imaging, followed by staining for DNA, EdU, NPAT and nascent histone RNA FISH (G) or pNPAT T1270 (H). (I) Representative images of cells treated with DMSO, 1µM APH, 500nM PF-07104091 (PF4091), or a combination of 1µM APH and 500nM PF-07104091 (PF4091). (J) Column 1: Raw single-cell data. Column 2: Average EdU signal and 95% confidence intervals as a function of time since anaphase for $CDK2^{inc}$ and $CDK2^{low}$ cells treated with DMSO, as well as $CDK2^{inc}$ cells treated with 1µM APH, 500nM PF-07104091 (PF4091), or a combination of 1µM APH and 500nM PF-07104091 (PF4091) (see Figure 5G).

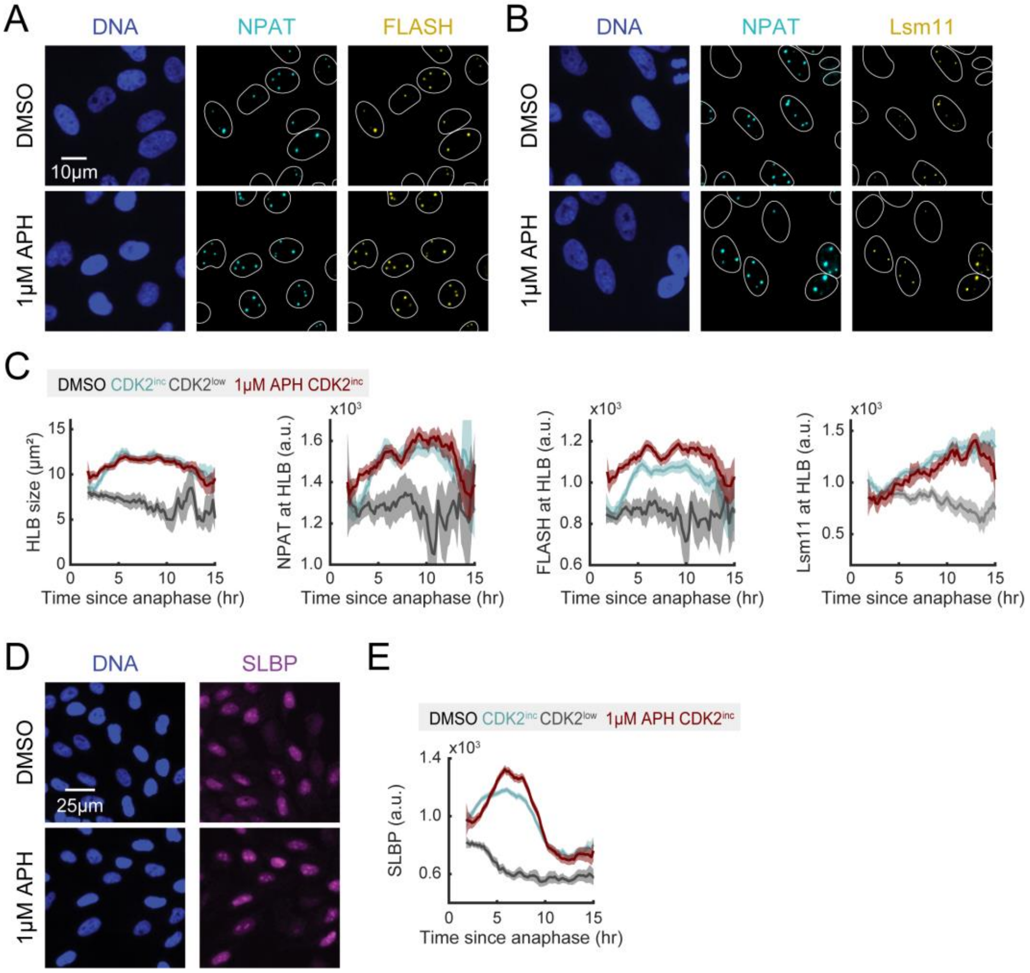


**Figure S7. HLB formation and SLBP levels upon DNA synthesis inhibition, related to Figure 5.** (A-B) Representative images of cells treated with DMSO or 1μM APH in the last 1 hr of imaging, followed by staining for DNA, NPAT, and FLASH (A) or Lsm11 (B). (C) Column 1-4: Average HLB size (1) and signal at the HLB for NPAT (2), FLASH (3), and Lsm11 (4) in CDK2$^{inc}$ and CDK2$^{low}$ cells treated with DMSO for 1 hr before fixation, with CDK2$^{inc}$ cells plotted for cells treated with 1μM APH for 1 hr before fixation. (D) Representative images of cells treated with DMSO or 1μM APH in the last 1 hr of imaging, followed by staining for SLBP. (E) Average nuclear signal for SLBP in CDK2$^{inc}$ and CDK2$^{low}$ cells treated with DMSO, and CDK2$^{inc}$ cells plotted for cells treated with 1μM APH for 1 hr before fixation.

| Gene | Probe sequence (reverse complement) | Position relative to the cleavage site | Gene | Probe sequence (reverse complement) | Position relative to the cleavage site |
|---|---|---|---|---|---|
| HIST1 | | | H3C11 | Agcttccttttggtgacaat | position 1 |
| H1-1 | Agctcttttcctgaaatgcg | position 1 | H3C12 | Cagctatcctcagtgagaat | position 2 |
| | Ggggatttcatctgtgtact | position 23 | | Aagacagtcgaccaatggtg | position 25 |
| H1-5 | Agccatttttagaggtctc | position 1 | H4C1 | Cagctcttcttttgtttagt | position 1 |
| | Cctgatctaacatgttgtgc | position 23 | H4C3 | Caacgcatttcagctcttta | position 11 |
| H1-2 | Ccagctcttttattgagatc | position 3 | H4C4 | Tactctttctcctgagtgga | position 1 |
| H1-3 | Cttcagctcttttgactgag | position 6 | | Ctacacttttagcagtgact | position 23 |
| H1-4 | Acagctcttttactgagagc | position 3 | H4C5 | Aagtcatgagctcttttgcg | position 9 |
| H2AC1 | Acgttttgtaacgtacagcc | position 15 | H4C6 | Cttcgtgtgacgaaggaggt | position 1 |
| H2AC4 | Tccaaagcacaacagctcat | position 13 | H4C7 | Gagcccttcggaagaaaacg | position 1 |
| H2AC6 | Tacaagcagtgaggttagct | position 16 | H4C11 | Accactcttcagctgagaaa | position 2 |
| H2AC7 | Atgtgagccccttttaaggaa | position 5 | H4C12 | Cggctcctcagctgagaaag | position 1 |
| H2AC8 | Gatttcaggcatcgtatgtg | position 25 | H4C13 | Tagtctacagctctttcaca | position 8 |
| H2AC12 | Tttctctaagacagtgtaga | position 2 | HIST2 | | |
| | Aacaaacatgactgtgtcca | position 24 | H2AC21 | Cagcccttcatttgacactt | position 1 |
| H2AC13 | Tgctcagttcttccacaaaa | position 4 | | Agctaatgcgctatttcgtg | position 23 |
| H2AC14 | Ggactcgtgtcatcttttta | position 8 | | Aaaggctctttttagagcca | position 62 |
| H2AC15 | Acacagctctacttggtgaa | position 5 | H2AC20 | Acgaaatagcgcattagctc | position 15 |
| H2AC16 | Cagctctcccgcagaaatag | position 1 | | Agtgtcaaatgaagggctga | position 37 |
| | Gttccaaatgtttaaccagt | position 25 | | Aaggctcttttaagagccac | position 60 |
| H2AC17 | Cagttcttccgtagaaatgg | position 1 | H2BC18 | Acagctgctttctcgattac | position 1 |
| | Taggtttggagaacagagtg | position 23 | | Tcaaacgctctgacaagtgt | position 23 |
| H2BC1 | Cacagctgtttcagtatgtt | position 1 | | Actccctacctgaccaaaag | position 46 |
| H2BC3 | Tatagctctccttgtgacaa | position 3 | | Cgaatgcatccgtgaccatg | position 71 |
| H2BC6 | Catgtcagtatatagctcca | position 13 | H2BC21 | Gctcttttctagtgattagg | position 1 |
| H2BC7 | Atgcaggtaattacaactct | position 14 | | Ccatgaccttcacactaaaa | position 63 |
| H2BC10 | Ccaatgccaagtctggaaaa | position 24 | | Aaaaagctacgtatgccatt | position 85 |
| H2BC12 | Gtttacagctctaatatcga | position 6 | H3C15 | Ggtacagcttcttccaagtg | position 7 |
| H2BC13 | Cccagtgataggaagagcga | position 24 | | Tctaatgccggaggagcacg | position 37 |
| H2BC14 | Agtgaactggctctttcttg | position 8 | | Cgttttgttgggagcctaag | position 75 |
| H2BC17 | Aaatgcacaggctctttctc | position 8 | H3C13 | Gtacagctgctcttgatgaa | position 7 |
| H3C1 | Ggtttcttacagctacttta | position 4 | | Gacgcacatcagatggagag | position 30 |
| H3C2 | Acagctactttcgttggaat | position 1 | | Tttgggttctgtaacgtcac | position 91 |
| H3C3 | Accgaatgggttacaactgt | position 16 | H4C14 | Caactcctcagatgagcaag | position 1 |
| H3C6 | Caaaggattacagccctttt | position 12 | | Tactgttgcactttacgcag | position 30 |
| H3C7 | Acggcgctttcaactgaaat | position 5 | | Ccaaagttaccttccaacat | position 53 |
| H3C8 | Cacagctattttcgggtgaa | position 3 | | Tttccgttaacttcagatcc | position 89 |
| H3C10 | Aaacagatcttttcagcgcg | position 5 | | Cgacttaagtctttacggct | position 133 |

Supplementary Table S1. Nascent histone mRNA probe set. Related to Key Resources Table.
